# Brain deviation scores derived from normative modelling identify dimensions of psychopathology in children and adolescents

**DOI:** 10.64898/2026.09.12.751102

**Authors:** Zheng Li, Bingqian Ren, Hao Wu, Zhidong Wang, Libin Zhang

## Abstract

Psychiatric diagnoses lack neurobiological grounding, and brain-based abnormalities correspond poorly to established diagnostic categories. While multivariate algorithms extract transdiagnostic dimensions from brain–behavior covariation, most studies rely on conventional structural morphometric measures such as cortical thickness and surface area, which fail to capture atypical developmental trajectories. Normative modeling (NM) quantifies individual-level brain deviation scores by contrasting brain structural features with population-derived percentiles, yielding continuous indices of atypical brain development. However, whether these scores can identify transdiagnostic dimensions remains unclear. We applied sparse canonical correlation analysis (sCCA) to investigate covariation between brain-deviation scores and psychiatric symptoms in 1095 children and adolescents from the Healthy Brain Network. We identified an emotion-dysregulation dimension (r = 0.33). When generalized to the independent cohort, this dimension showed nominal significance (r = 0.18, *P*_FDR_ = 0.05, uncorrected P = 0.01, n = 198). By contrast, no significant dimension was detected using conventional structural brain features. Emotion dysregulation was primarily driven by deviations in cortical thickness and surface area, with distinct regional distributions; reductions in cortical surface area in the middle frontal gyrus, cingulate gyrus, superior temporal gyrus, and insula carried the highest weights. Latent brain scores for emotion dysregulation, computed using sCCA weights for brain deviation scores, showed specific correlations with out-of-model emotion dysregulation-related phenotypes. In addition, females exhibited significantly higher latent brain scores than males. Collectively, NM-based brain deviation scores were more sensitive than conventional structural metrics in identifying transdiagnostic psychiatric dimensions. Furthermore, latent brain scores derived from sCCA-estimated weights may serve as a straightforward metric that captures the degree of dimension-specific atypical brain deviation.

## INTRODUCTION

Childhood and adolescence represent a critical window for brain maturation, as well as the peak period for onset of many psychiatric disorders ^1–3^. Reliable biomarkers for pediatric psychiatric disorders are therefore essential for early identification, diagnosis, and intervention. However, accumulating evidence indicates that categorical diagnostic frameworks have limited biomarker discovery efforts ^4^. These categorical systems fail to adequately account for widespread comorbidity and symptom overlap, as well as divergent clinical profiles among patients receiving the same diagnosis ^5–7^.

Accordingly, there is growing consensus that research should move beyond categorical diagnostic systems toward dimensional classifications of psychopathology. The Hierarchical Taxonomy of Psychopathology (HiTOP) represents one of the most prominent dimensional frameworks ^8^. The HiTOP model hierarchically organizes behavioral symptoms to identify shared and unique latent components across psychiatric conditions, which accounts for widespread comorbidity and variable clinical presentations. Nevertheless, HiTOP dimensions are defined based on clinical behavioral assessments yet lack underlying neurobiological foundations ^5^.

A growing body of work argues that dimensional models of psychopathology should not rely solely on behavioral symptoms but also incorporate neuroimaging information ^9,10^. Frameworks integrating neuroimaging and behavioral measures can harness the respective strengths of both modalities: neuroimaging signatures serve as candidate neural markers for behavioral presentations, whereas behavioral data lend clinical interpretability to neuroimaging findings. Multivariate machine learning approaches are well-suited for such analytical frameworks, as they jointly model complex brain– behavior associations by optimizing feature weights to maximize linear correlation between neural and behavioral measures ^11^.

Structural brain morphometric measures have been widely applied to define dimensions of psychopathology. One study using the ABCD dataset reported that the general psychopathology (p-factor) cluster, internalizing symptom cluster, and neurodevelopmental symptom cluster were associated with alterations in cortical thickness and surface area across widespread brain regions ^12^. Another investigation drawing on the IMAGEN dataset identified associations between the excitability-impulsivity symptom cluster and surface area alterations in the dorsolateral prefrontal cortex, anterior cingulate cortex, and inferior parietal cortex ^9^. However, most of these studies rely on conventional structural brain metrics, which only capture physical brain morphology and cannot assess whether observed structural features fit age-typical developmental patterns. This limitation is comparable to measuring children’s height and weight without reference to age-matched normative ranges.

Normative modeling (NM) represents a statistical framework for characterizing individual-level deviations from normative brain development ^13^. Leveraging large-sample normative cohorts, NM models the mean and inter-individual variability of neuroimaging feature distributions as nonlinear functions of age and other covariates (e.g., sex, intracranial volume, scan site). Using this normative reference, each individual’s neuroimaging features are compared against the age-matched distribution to compute individualized brain deviation scores. Such brain deviation scores have been applied to the study of autism spectrum disorder ^14^, attention-deficit/hyperactivity disorder ^15^, depression and anxiety ^16,17^. Despite these advances, to our knowledge, no study has yet delineated psychopathological dimensions using multivariate algorithms based on brain-deviation scores.

Compared with structural brain features, brain deviation scores reflect deviations from normative developmental trajectories. Accumulating evidence indicates that deviation scores outperform structural brain features across multiple clinical populations and outcomes. Studies among children and adolescents have shown that brain deviation scores outperform structural features in predicting psychopathology dimensions ^18^. Evidence from adult samples further indicates that these deviation scores yield higher predictive performance for cognitive outcomes ^19^. Furthermore, these deviation scores achieve superior classification accuracy for schizophrenia and early psychosis ^20^. Compared with structural features, brain deviation scores also produce larger effect sizes for group differences in schizophrenia ^19^ and lower false-positive rates for bipolar disorder under small-sample conditions ^21^.

Previous studies have demonstrated that the associations between regional wise brain deviation scores and psychopathology vary across brain regions and are modulated by age and sex ^22,23^. Consequently, whole-brain region wise deviation scores are not readily interpretable for clinicians. One key advantage of multivariate algorithms is the generation of latent scores from linear combinations of multiple input variables. This approach condenses high-dimensional data to facilitate interpretability and yields potential phenotypic metrics for assessing brain behavioral abnormalities in psychiatric disorders ^24,25^. By identifying covariation between brain and behavioral measures, multivariate algorithms assign weights to individual brain deviation scores to compute aggregated latent scores. This provides a promising avenue for improving the interpretability of brain deviation scores.

Despite growing interest in neuroimaging-informed delineation of psychopathological dimensions, existing studies have been limited in several respects. First, it remains unknown whether NM-derived brain deviation scores can be used to delineate psychopathological dimensions in children and adolescents. Second, it is unclear whether brain deviation scores offer greater sensitivity for identifying psychopathological dimensions relative to structural brain features. Finally, it remains unclear whether latent brain scores derived from the brain deviations can reflect dimension-specific psychopathological symptoms.

The present study seeks to address these unresolved questions. We adopted a multivariate algorithm to identify covariation between brain deviation scores and psychopathology, with the aim of delineating psychopathological dimensions and their dimension-specific brain deviation patterns. We hypothesized that brain deviation scores derived from normative modelling would exhibit higher sensitivity for parsing psychopathological dimensions compared with conventional raw brain structural features. Furthermore, latent brain scores computed from the weights of the multivariate algorithm could reflect dimension-specific patterns of brain abnormalities.

## MATERIALS AND METHODS

### Participants

The discovery dataset was derived from the Healthy Brain Network (HBN), a large-scale multimodal cohort of children and adolescents aged 5 to 21 years recruited in New York City, USA ^26^. The final analytical sample comprised 1,095 participants. The mean age of participants was 11.14 ± 3.41 years (range: 5.06–21.9 years). The sample included 406 females (37%) and 689 males (63%). Of these participants, 434 (40%) were typically developing, whereas 661 (60%) had received at least one psychiatric diagnosis (Supplementary Figure 1; Supplementary Table 1).

The independent dataset was derived from the NKI-Rockland Sample (NKI-RS), a longitudinal cohort of individuals aged 6 to 18 years that focuses on child and adolescent populations ^27^. Baseline data from the NKI-RS cohort were utilized for generalization analyses, as this time point had the largest available sample compared with other time points. The final sample for generalization analyses comprised 198 participants. Participants had a mean age of 12.9 ± 3.1 years (range: 6.17–17.92 years). The sample consisted of 93 females (47%) and 105 males (53%). Among these participants, 114 (58%) were typically-developing and 82 (42%) had at least one psychiatric diagnosis. Two NKI-RS participants lacked diagnostic data, yielding 196 participants with complete diagnostic and comorbidity information (Supplementary Figure 1; Supplementary Table 2).

Diagnostic procedures for mental disorders in the HBN and NKI-RS datasets are described in Supplementary Methods.

### Behavioral assessment

In discovery and independent cohorts, The parent-reported Child Behavior Checklist (CBCL) ^28^ were used to examine brain-deviation associations (Figure 1B). The 119-item CBCL evaluates children’s mental-health difficulties over six months with a 3-point Likert scale. During preprocessing, items with more than 1% missing data were excluded, leaving 117 valid items for analysis (Supplementary Table 3). Item scores were residualized for age and sex using linear regression.

**Figure 1.**
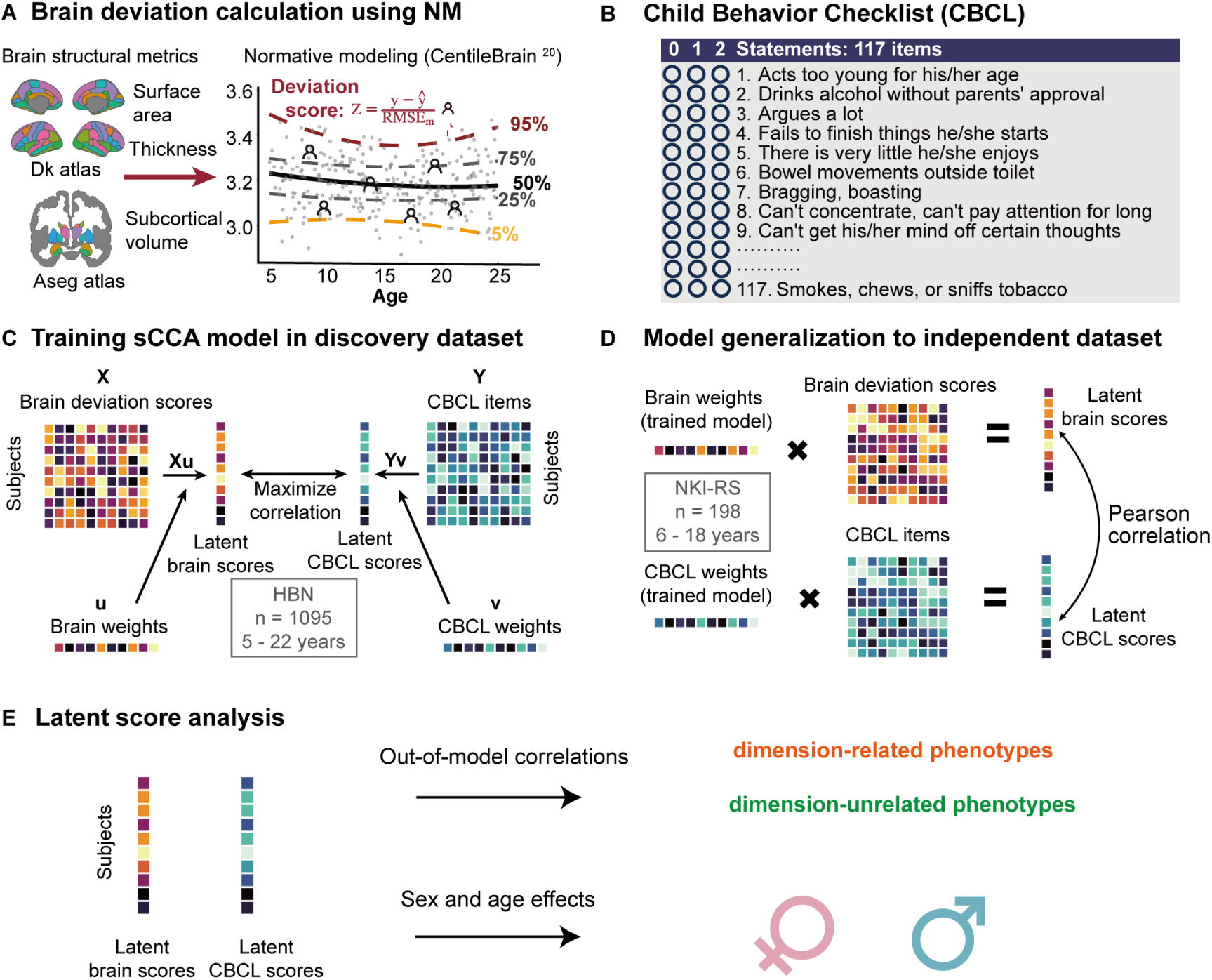
Schematic overview of the data-analysis workflow. (**A**) Calculation of brain deviation scores. NM was applied to compute brain-deviation scores for cortical thickness, surface area, and subcortical volumes. (**B**) Symptom assessment. Psychopathological symptoms were assessed using the Child Behavior Checklist (CBCL) in both discovery and independent datasets. (**C**) sCCA model training. Sparse canonical correlation analysis (sCCA) was trained on brain-deviation scores and CBCL symptom items within the discovery dataset. (**D**) Model generalization to the independent dataset. Pre-trained sCCA weights from the discovery dataset were applied to the independent dataset to derive latent brain and CBCL scores, whose associations were quantified via Pearson correlation. (**E**) Latent-score analyses. Partial correlations between sCCA-derived latent scores and phenotypes that were not included in sCCA modeling. We further examined age- and sex-related effects on these latent scores.

For generalizability validation, we used the identical set of 117 residualized CBCL item scores in the independent cohort to ensure consistent input variables across datasets.

### MRI acquisition

T1-weighted structural MRI (sMRI) scans for the HBN cohort were acquired across four sites. The Staten Island (SI) site used a 1.5 T Siemens Avanto scanner (TR = 2730 ms, TE = 1.64 ms, TI = 1000 ms, flip angle = 7°, 176 slices, 1.0 mm isotropic voxels). The Rutgers University Brain Imaging Center (RUBIC) deployed a 3.0 T Siemens Tim Trio scanner (TR = 2500 ms, TE = 3.15 ms, TI = 1060 ms, flip angle = 8°, 224 slices, 0.8 mm isotropic voxels). The CitiGroup Cornell Brain Imaging Center (CBIC) and CUNY shared identical 3.0 T Siemens Prisma hardware and used the same pulse-sequence parameters as RUBIC.

For the NKI-RS cohort, all T1-weighted sMRI data were acquired using a 3.0 T Siemens Tim Trio scanner at the Nathan Kline Institute. The acquisition sequence parameters were set as follows: TR = 1,900 ms, TE = 2.52 ms, TI = 900 ms, flip angle = 9°, 176 slices, and 1.0 × 1.0 × 1.0 mm isotropic voxel resolution.

Complete imaging protocols can be retrieved (HBN: https://fcon_1000.projects.nitrc.org/indi/cmi_healthy_brain_network/MRI_Protocol.html. ; NKI-RS: http://fcon_1000.projects.nitrc.org/indi/enhanced/mri_protocol.html).

### Quality control and processing of sMRI data

To avoid cross-cohort methodological discrepancies, all sMRI data from discovery and validation cohorts underwent unified quality control and standardized preprocessing implemented by the Reproducible Brain Charts consortium ^29^, including visual scan assessment, image correction, skull stripping, brain reconstruction, and atlas-based feature segmentation; detailed processing procedures and extracted structural features are provided in the Supplementary Materials.

Cortical thickness and surface area were quantified based on the Desikan–Killiany (DK) ^30^ atlas across 68 bilateral cortical regions, generating 136 cortical features (68 regions × 2 morphological metrics). Subcortical volumes were segmented using the automatic segmentation (ASEG) ^31^ atlas, yielding 14 subcortical volumetric features, with a total of 150 extracted structural brain features (Supplementary Table 4).

To remove scanner-related site batch effects in multi-center data, structural brain metrics were harmonized via the neuroHarmonize toolbox ^32^, which implements the ComBat-GAM procedure. Age was included as a nonlinear smooth term to capture continuous developmental variation. The harmonized structural features from the discovery dataset were then fed into the NM model to compute brain deviation scores. Both these deviation scores and the harmonized structural features were subsequently used to construct the sCCA model.

### Computation of individual brain deviation scores via normative modeling

After obtaining brain-structural features for each participant, we used CentileBrain to compute individual brain deviation scores (Figure 1A). CentileBrain is a publicly available NM for brain structural morphology ^20^. This model was trained on regional morphometric data from 37,407 healthy individuals aged 3 to 90 years, aggregated from 87 multi-ethnic global datasets. CentileBrain were constructed separately for males and females. Previous validations have verified that CentileBrain achieves high predictive accuracy across the entire lifespan, maintains stable longitudinal performance over a 2-year interval and yields robust normative estimation across diverse ethnic populations ^20^.

The regional brain deviations (namely Z-scores) were calculated via the formula below:

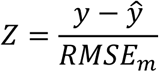

where y refers to the observed brain morphological value, *ŷ* denotes the normative predicted value from NM, and *RMSE_m_* is the root mean square error of NM. Positive Z-scores indicate brain morphological metrics higher than age-matched normative predictions, while negative Z-scores indicate lower values.

### Sparse canonical correlation analysis between brain deviations and psychiatric symptoms

As illustrated in Figure 1C, we adopted sparse canonical correlation analysis (sCCA) to identify covariation patterns between CBCL items and brain deviation scores using the R package PMA (v. 1.2-4) ^33^. Through sparsity regularization, sCCA enhances model interpretability and lowers the risk of overfitting in brain-behavior association analyses. See Supplementary Methods for details.

Let ***X****_n_*_×*p*_(brain-deviation features) and ***Y****_n_*_×*q*_(clinical-symptom features) represent feature matrices for *n* participants, with *p* brain features and *q* clinical-symptom indicators. Brain deviation scores and CBCL item scores were standardized across participants to equalize variances. Per Equation (1), sCCA optimizes loading vectors **u** and **v** to maximize canonical correlation between X**u** and *Y***v**. Unit *L*_2_-norm constraints on **u** and **v** to ensure scale identifiability, while distinct *L*_1_ sparsity parameters *c*_1_ and *c*_2_ impose sparsity on **u** and **v**.

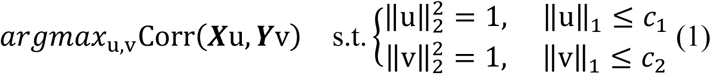

The sCCA sparsity parameters *c*_1_ and *c*_2_ were optimized via grid search. Parameter values were chosen to maximize average canonical correlation across repeated resampling runs.

Permutation testing evaluated the statistical significance of each canonical variate. With the brain-deviation matrix kept fixed, row indices of the clinical-symptom matrix were permuted, and sCCA was re-run 1000 times using identical regularization parameters to build the null correlation distribution.

We used a bootstrap resampling framework with 1000 iterations to quantify feature stability within each canonical component. At each iteration, two-thirds of participants from the discovery cohort were randomly subsampled; the remaining one-third of observations were generated by resampling with replacement from this subset. Feature-weight distributions across resampling runs were used to construct corresponding confidence intervals. A feature was defined as stable when its resampling-based 95 % confidence interval (CI) excluded zero. Given their high dimensionality, brain-deviation features adopted a stricter 99 % CI threshold.

### Comparison of sCCA models using brain deviation scores versus conventional brain structural features

To compare how well brain deviation scores and conventional structural features delineate psychopathological dimensions, we applied the identical pipeline for multivariate brain-behavior analyses. Only the input brain metrics differed: conventional brain structural features were used instead of brain deviation scores derived from these measures. We then contrasted the two sets of sCCA models on explained variance and the identification of the canonical variates.

### Model generalization analysis

To assess generalizability, we applied the sCCA loading weights (**u** for brain, **v** for behavioral) trained on the discovery dataset to the independent dataset. Generalizability was determined by Pearson’s correlation between latent brain and behavioral scores; a canonical variable was deemed generalizable if this cross-cohort correlation reached significance after FDR correction (Figure 1D).

### Specificity of associations between latent scores and out-of-model phenotypes

Four distinct psychopathology dimensions (emotion dysregulation, externalizing behavior, eating pathology, and substance-use-psychosis) were first identified in the discovery dataset. When tested in an independent cohort, only the emotion dysregulation dimension held across cohorts (see Results 3.1). To test the specificity of emotion-dysregulation latent scores (brain-side and behavior-side separately), we calculated partial correlations between latent scores of all four dimensions and out-of-model phenotypes in the discovery dataset, partialing out variance from the other three dimensions for each analysis. We hypothesized that if emotion-dysregulation latent scores reflected true specificity, these scores would show strong correlations with emotion-dysregulation-related phenotypes and weak correlations with non-emotion-dysregulation phenotypes (Figure 1E).

Phenotypes were selected to distinguish emotion dysregulation-related outcomes from comparison phenotypes. Specifically, irritability was selected as an emotion-dysregulation-related phenotype given its status as a core feature of emotion dysregulation ^34–36^. Callous-unemotional traits, body mass index (BMI), and cocaine-related substance involvement were included as non-emotion-dysregulation comparison phenotypes, given their documented associations with the three other dimensions identified in the present study ^37–39^. Detailed measurement of each phenotype is provided in Supplementary Methods.

### Analysis of age effects and sex differences

Using the discovery dataset, we further tested for sex and age effects in the latent scores of emotion dysregulation (Figure 1E). Specifically, latent brain and behavioral scores for emotion dysregulation were computed by multiplying standardized brain and behavioral data by their corresponding dimension weights. Ordinary least-squares (OLS) linear regression models were then fitted to assess main effects of age, sex, and the age-by-sex interaction. For significant interaction terms, simple slope analysis was used to evaluate age-outcome associations separately in male and female subgroups.

## RESULTS

### Delineating psychopathology dimensions informed by brain deviation scores

Using the discovery dataset, we applied sCCA to detect multivariate patterns linking brain deviation scores to CBCL items. The top four canonical variates were retained according to the explained covariance (Figure 2A). These canonical variates collectively explained 30% of the variance linking brain deviations and psychopathology symptoms (Supplementary Table 5). As shown in Figure 2B, all four canonical variates reached statistical significance following permutation testing (canonical variate 1: *r* = 0.33, *p*_FDR_ < 0.01; canonical variate 2: *r* = 0.32, *p*_FDR_ < 0.05; canonical variate 3: *r* = 0.33, *p*_FDR_ < 0.01; canonical variate 4: *r* = 0.34, *p*_FDR_ < 0.001, Supplementary Figure 2).

**Figure 2.**
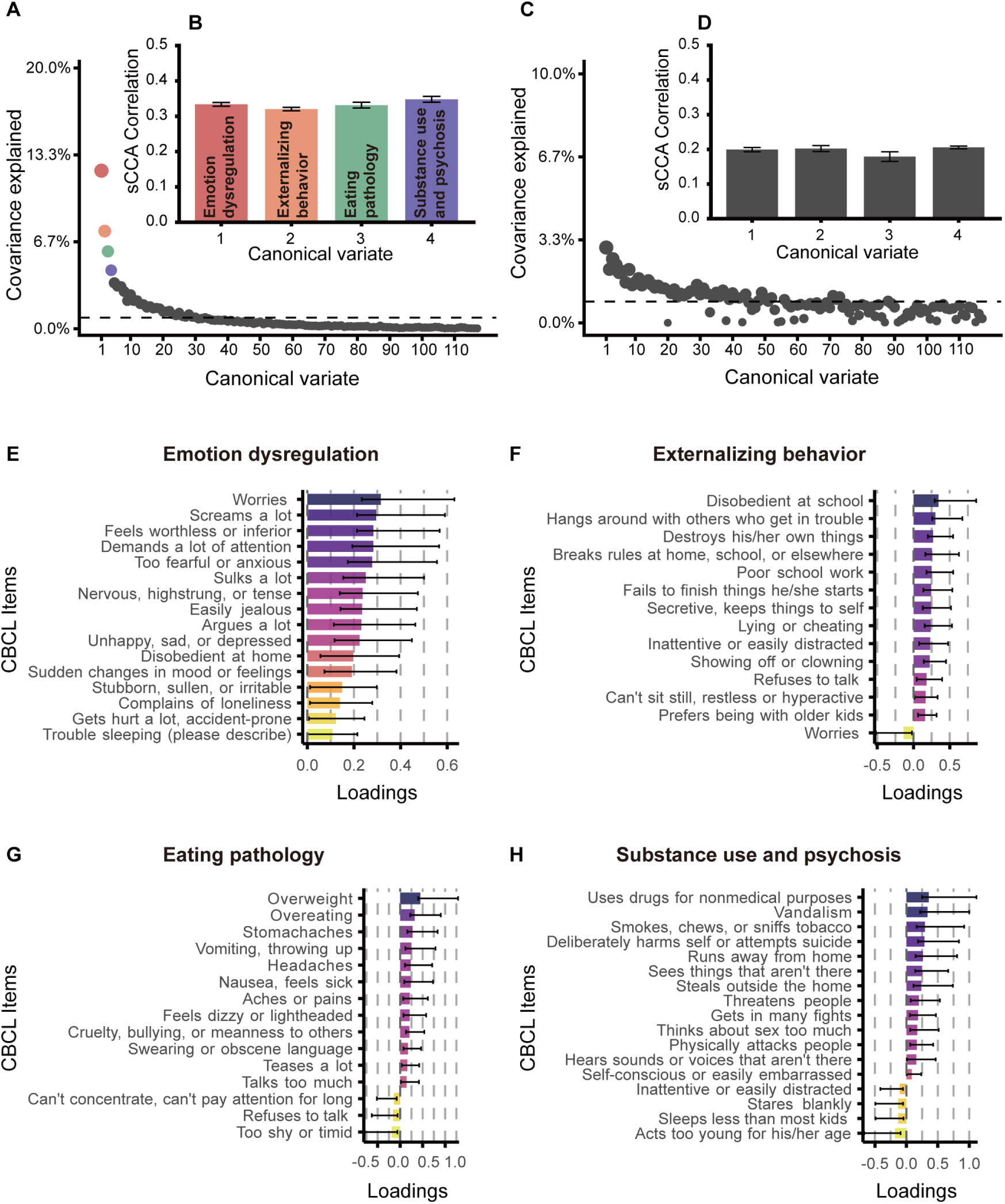
Multivariate brain-CBCL associations derived in the discovery dataset. (**A**) Variance explained by the brain deviation-based sCCA model; dashed line indicates the mean variance explained. (**B**) Latent brain–behavior correlation for the top four dimensions of the brain deviation-based sCCA model. Error bars denote the standard error from 10 resampling iterations. (**C**) Variance explained by structural the brain-based sCCA model. (**D**) Latent brain–behavior correlation for the top four dimensions of the structural brain-based sCCA model. (**E–H)** CBCL items with stable loadings on the psychopathological dimensions. Each bar shows the 95% CI of feature-weight distributions from bootstrapped resampling.

As shown in Figure 2E-H, canonical variate 1 was characterized by anxiety, aggressive behavior, and sudden mood shifts, and we termed this variate “emotional dysregulation”. Canonical variate 2 featured rule-breaking behavior, aggressive behavior, and attention problems and was termed “externalizing behavior”. Canonical variate 3 reflected overweight and binge-eating behavior and was labeled “eating pathology”. Canonical variate 4 captured non-medical drug use and psychotic-like behavior and was termed “substance use and psychosis”.

### Comparison of sCCA models: brain deviation scores versus conventional structural features

We applied the same pipeline to identify multivariate association patterns between conventional structural features and CBCL items. As shown in the scree plot (Figure 2C), no clear elbow for variance explained was observed. To maintain consistency with the brain deviation-based model, we also retained the top four canonical variables. These four components collectively accounted for 10% of the variance linking structural brain features to psychopathological symptoms (Supplementary Table6). Permutation tests revealed no statistically significant canonical components (Figure 2D).

### Generalization of multivariate brain–behavior association patterns in an independent sample

To verify the generalization of the model, we applied the brain and behavioral weights identified in the discovery dataset to the independent dataset. As shown in Figure 3, we observed a nominally significant association between latent brain and behavioral scores for the first canonical variate (r = 0.18, *P*_FDR_ = 0.05, uncorrected P = 0.01, permutation P = 0.01; see Supplementary Figure 3), whereas no significant correlation was observed for the second, third, and fourth dimensions.

**Figure 3.**
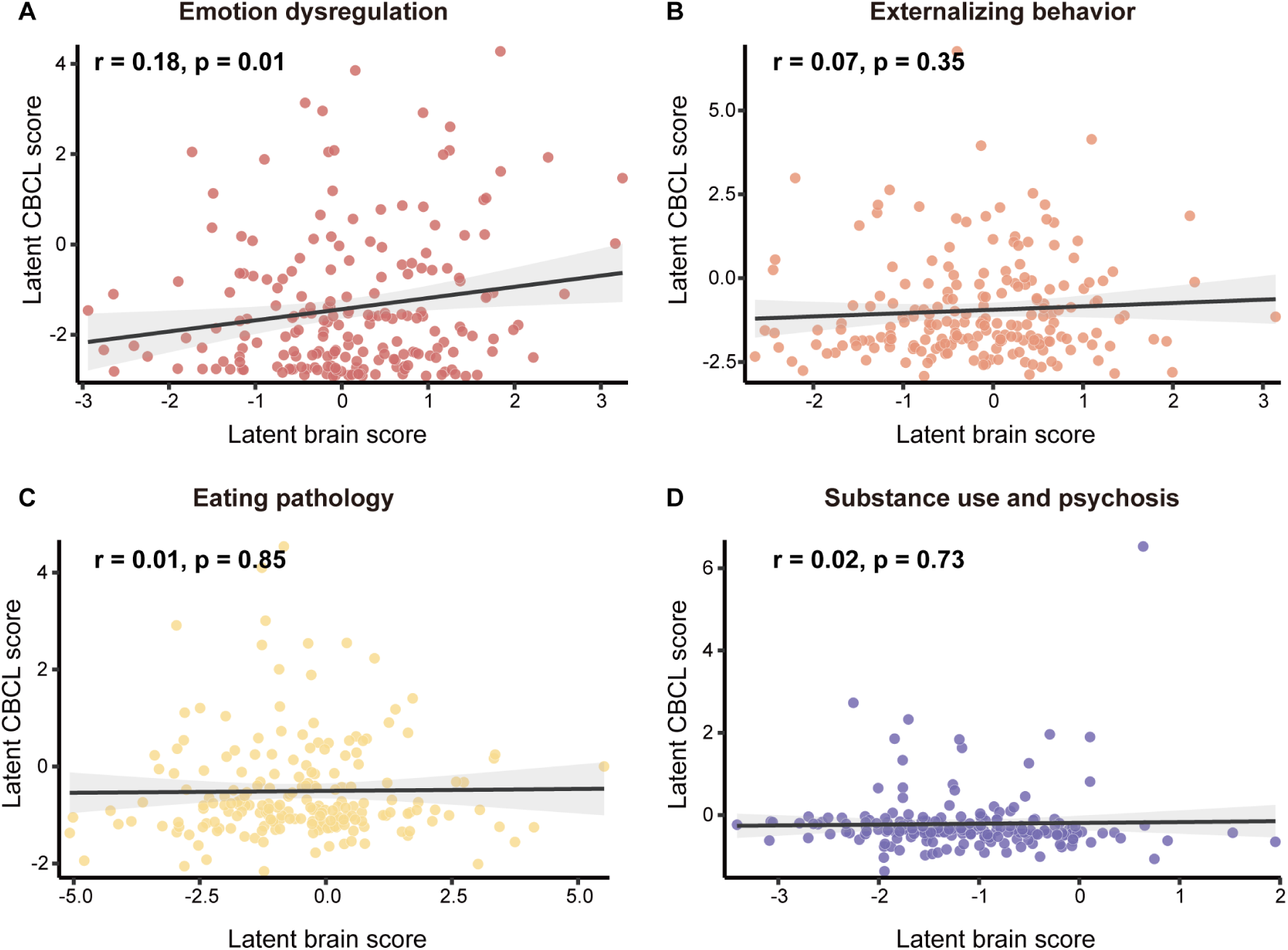
Generalizability analysis of canonical variates in the independent dataset. (**A–D**) Pearson correlations between latent brain scores and latent behavioral scores for the different psychopathology.

### The brain deviation pattern of psychopathology

We found that the emotion dysregulation dimension was associated with brain deviations in cortical surface area and cortical thickness. As shown in Figure 4, greater emotion dysregulation was associated with reduced surface area in the cingulate cortex, superior temporal cortex, insula, middle frontal gyrus, orbitofrontal cortex, inferior frontal cortex, inferior parietal lobule, and lingual gyrus. Additionally, more severe emotion dysregulation was associated with greater cortical thickness in the parietal lobe, precuneus, isthmus cingulate, and temporal cortex, as well as reduced cortical thickness in the fusiform gyrus, anterior cingulate cortex, paracentral lobule, and transverse temporal cortex. Subcortical volume deviations did not contribute significantly to the emotion dysregulation dimension. The brain deviation patterns for externalizing behavior, eating pathology, and substance use and psychosis are presented in the Supplementary Figure 4.

**Figure 4.**
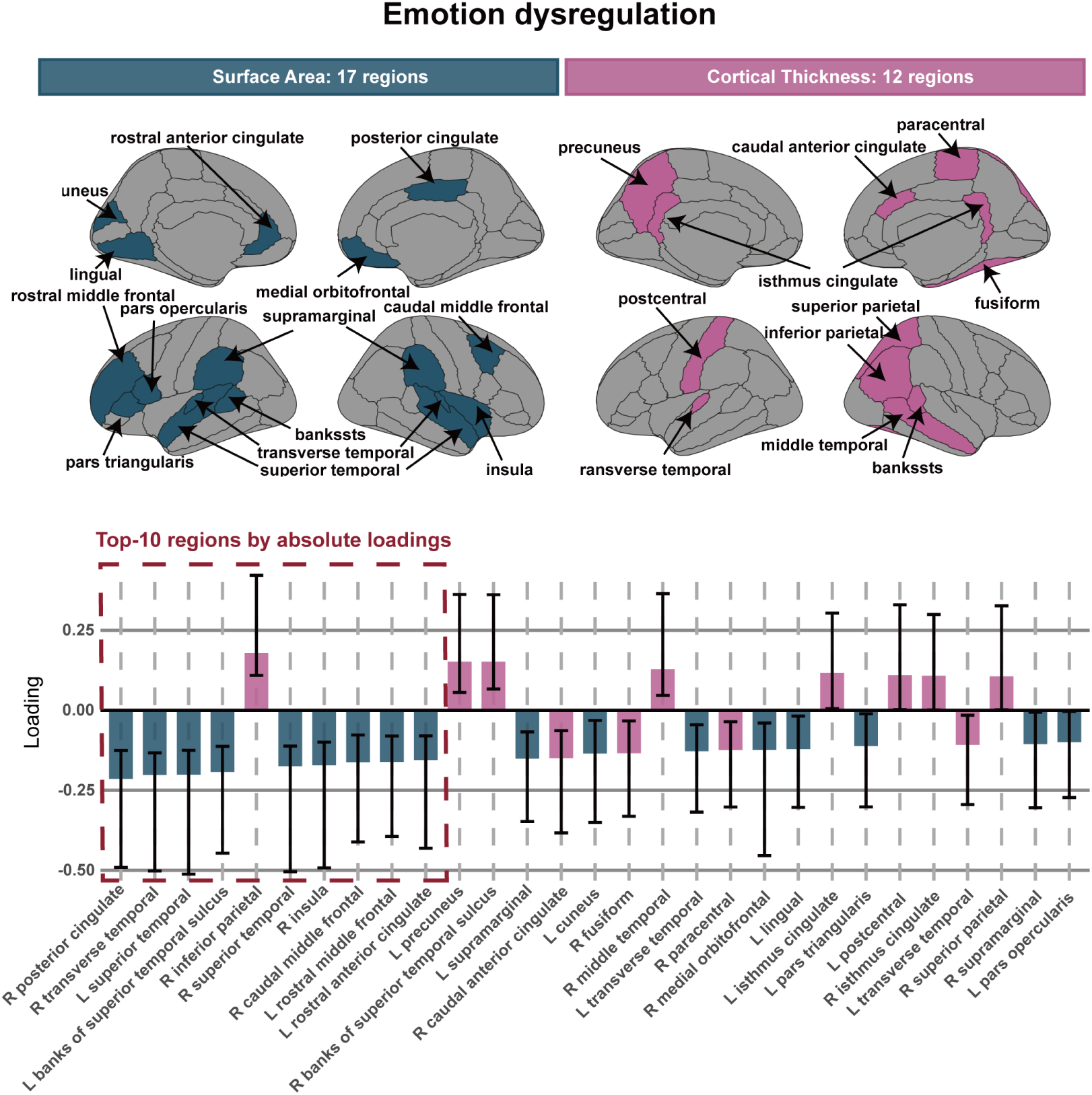
Brain deviation pattern of emotion dysregulation. The brain maps illustrate the spatial locations of stable contributing regions after resampling procedures. The bar plot displays loadings of each brain region along the emotion dysregulation dimension, sorted by absolute loading magnitudes in descending order from left to right. Each bar shows the 99% CI of feature-weight distributions from bootstrapped resampling. bankssts: banks of the superior temporal sulcus, L: left, R: right

Cortical surface area contributed more strongly to the emotion-dysregulation dimension compared with cortical thickness. When brain regions were ranked by the absolute magnitude of their weights, nine out of the top-ten regions showed contributions from surface area deviation (Figure 4), mainly distributed across the cingulate cortex, temporal lobe, insula, and prefrontal lobe. Furthermore, among regions with high contributions to the emotion-dysregulation dimension, 17 regions (59%) exhibited surface-related deviations, whereas 12 regions (41%) were associated with cortical-thickness metrics.

### Specificity validation of latent scores from psychopathological dimensions

We further examined the specificity of latent scores derived from the emotion dysregulation dimension.

Compared with the latent behavioral scores of the other three psychopathological dimensions, the latent behavioral score of the emotion dysregulation dimension showed the strongest associations with irritability (*r* = 0.491, *p*_FDR_ < 0.001) (Figure 5A left panel). By contrast, it exhibited weak correlations with callous-unemotional traits (*r* = 0.075, *p*_FDR_ < 0.05) and BMI (*r* = −0.102, *p*_FDR_ < 0.01), and no significant correlation was observed for cocaine-related substance involvement (Figure 5C-D, left panel).

**Figure 5.**
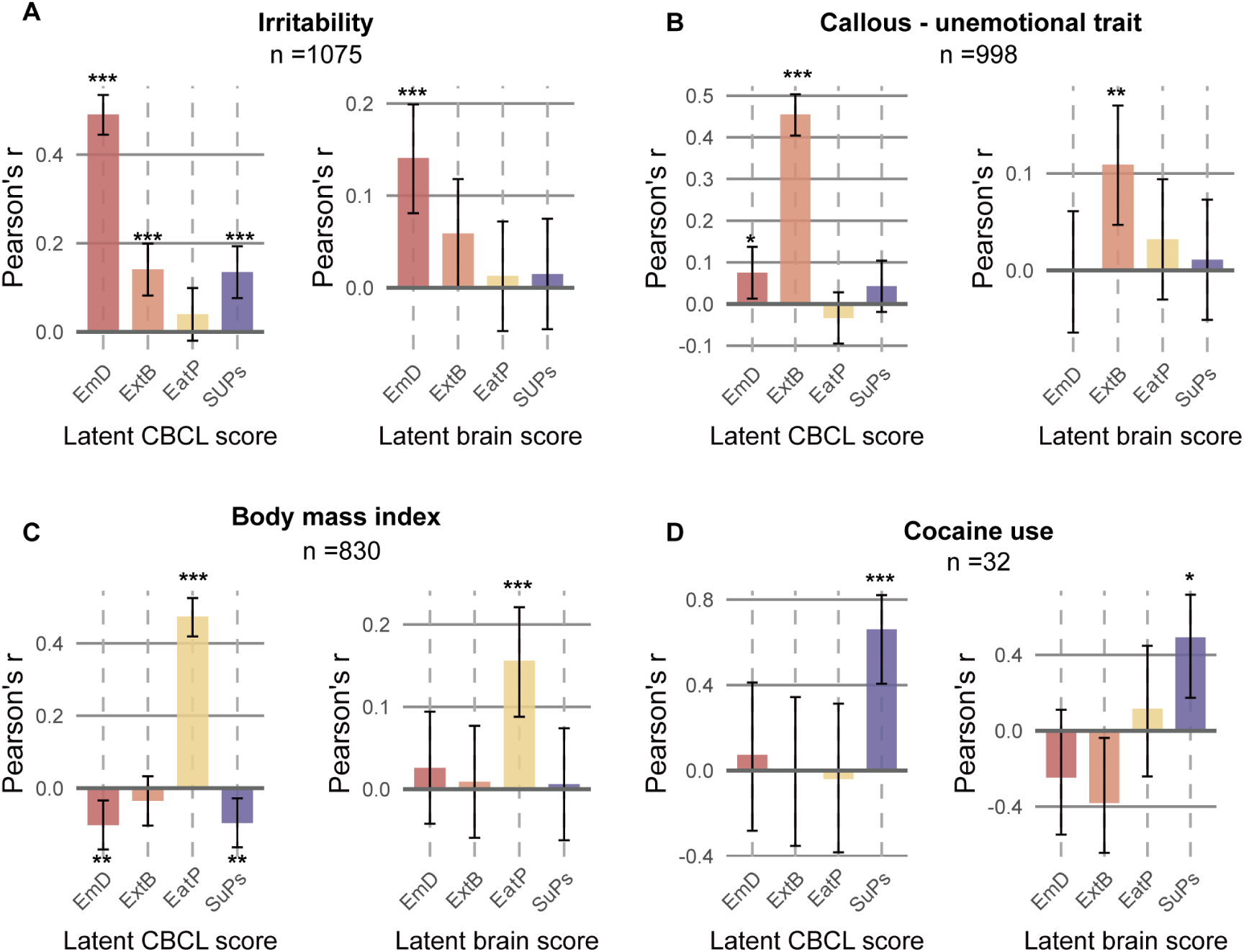
Associations between latent scores and independent behavioral phenotypes in the discovery dataset. (**A-D**) Partial correlations of canonical latent behavioral scores (left panel) and canonical latent brain scores (right panel) with out-of-model phenotypes. Error bars represent 95% confidence intervals. Owing to varying completion rates for different scales, sample sizes differ across measures; exact sample sizes are labelled in the figure. *** *P*_FDR_ < 0.001, \*\**P*_FDR_ < 0.01, * *P*_FDR_ < 0.05, † *P*_uncorrected_ = 0.04. EmD: emotion dysregulation, ExtB: externalizing behavior, EatP: eating pathology, SuPs: Substance use and psychosis.

Compared with the latent brain scores of the other three psychopathological dimensions, the latent brain score of the emotion dysregulation dimension showed the strongest associations with irritability (*r* = 0.141, *p*_FDR_ < 0.001) (Figure 5A-C right panel). By contrast, no significant correlations were observed for callous-unemotional traits, BMI, and cocaine-related substance involvement (Figure 5C-D, right panel).

### Age and sex effects and their interaction on dimension-specific latent scores

We further examined sex differences and age-related effects on latent scores of emotion dysregulation within the discovery dataset.

The emotional dysregulation latent behavioral score yielded a significant age-by-sex interaction (Figure 6A; *F*(1,1091) = 12.30, *p* < 0.001; adjusted*R*^2^ = 0.008). Simple-slope decomposition revealed divergent age trajectories: CBCL symptom scores declined with age in males (*b* = −0.056, 95%CI[−0.105, −0.006]) yet increased with age in females (*b* = 0.086, 95%CI[0.024,0.147]).

**Figure 6.**
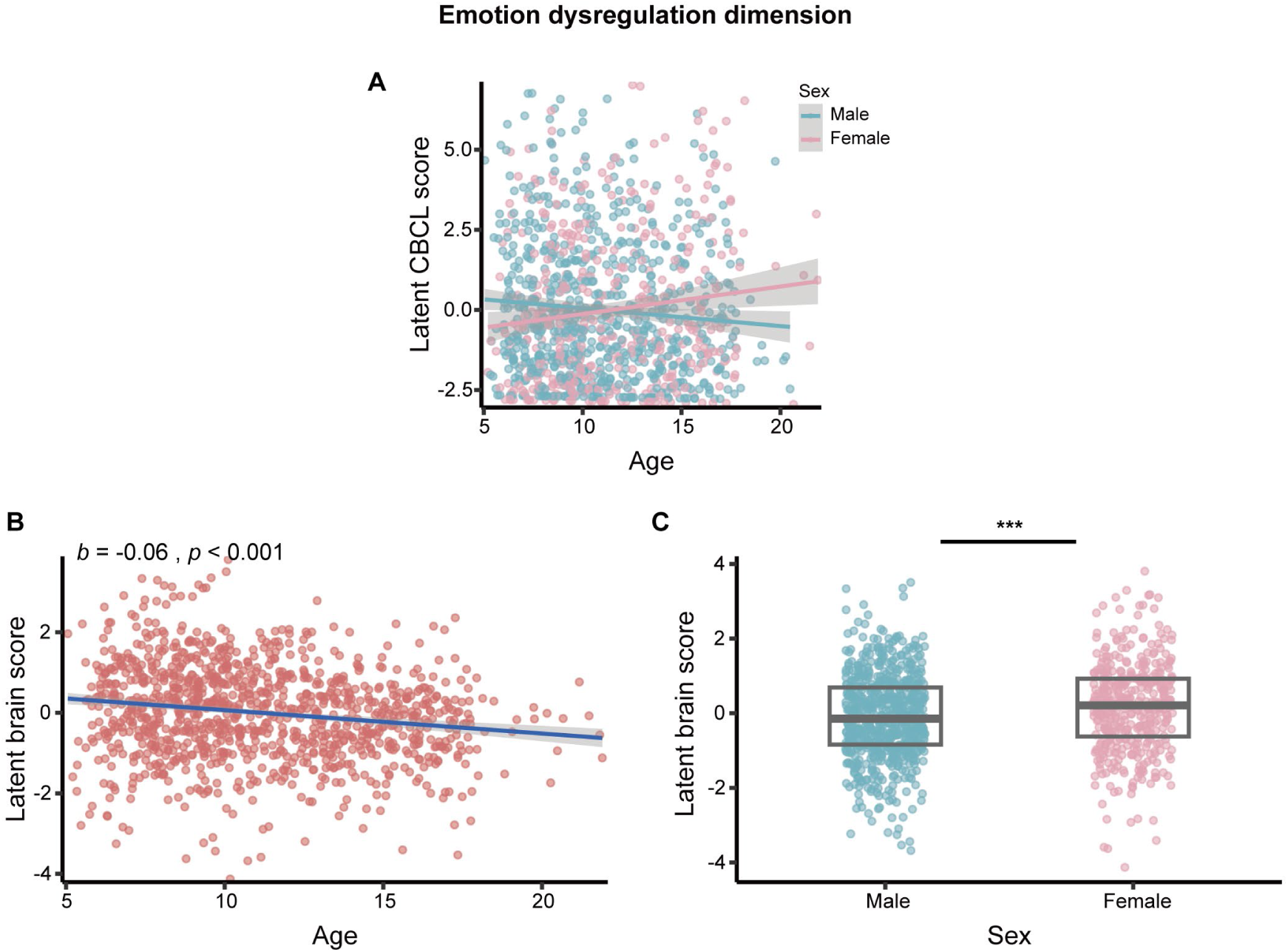
Age and sex effects on latent scores. (**A**) Age-by-sex interaction for behavioral latent scores. (**B**) Scatter plot of age versus brain latent score. The solid line denotes the regression fit, with gray shading indicating the 95 % confidence interval; each dot corresponds to one participant. (**C)** Box-and-scatter plot illustrating latent brain score distributions stratified by sex. Overlaid points show individual data; box edges mark the interquartile range (25th and 75th percentiles), and the central line denotes the median. *** *p* < 0.001.

For the latent brain score (Figure 6B-C), significant main effects of age (*F*(1,1092) = 35.84, *p* < 0.001) and sex (*F*(1,1092) = 15.19, *p* < 0.001) were observed. Brain-deviation scores decreased as age advanced (*b* = −0.061, 95%CI[−0.081, −0.041]). Age-adjusted estimated marginal means showed that females had significantly higher latent brain scores than males (*p* < 0.001, adjusted Cohen’s *d* = 0.25). The age-by-sex interaction was not statistically significant for brain latent scores. In summary, the emotion dysregulation brain-deviation pattern decreased with age across both sexes, with females exhibiting higher scores.

## DISCUSSION

We applied sCCA to map multivariate associations between brain-deviation scores and behavioral phenotypes for deriving psychopathological dimensions. We identified an emotion-dysregulation dimension in the discovery dataset and, critically, this dimension yielded a nominally significant correlation in the independent cohort. By contrast, no significant brain-behavior associations emerged for conventional structural brain features using identical data and analytical pipelines. Further analyses revealed that the emotion dysregulation dimension was linked to deviations in cortical thickness and surface area with divergent spatial distribution patterns. Surface area reductions within the prefrontal, superior temporal, cingulate, and insular regions carried the highest weights for this dimension. Latent brain scores of emotion dysregulation showed specific associations with out-of-model phenotypes. In addition, these latent brain scores also showed age- and sex-related effects.

Compared with sCCA trained on conventional brain structural features, sCCA using brain deviation scores explained higher variance and yielded significant psychopathological dimensions. This finding indicates that brain deviation scores exhibit higher sensitivity for delineating psychopathological dimensions relative to conventional brain structural features, which is consistent with previous evidence. One study demonstrated that brain deviation scores outperformed gray matter volume in predicting transdiagnostic dimensions^18^. Extending this prior work, the present study adopted multivariate algorithms and further confirmed that brain deviation scores, rather than conventional brain structural features, confer greater sensitivity for identifying psychopathological dimensions.

Multimodal data fusion is a key direction for neuroimaging-informed transdiagnostic dimensions^9,12^. Lett et al. combined task-based activation, brain structural features, and white matter metrics. Relative to earlier studies using fewer imaging modalities, their work identified broader transdiagnostic dimensions^9^. One dimension labeled “emotional and behavioral dysregulation” resembled the emotion dysregulation dimension derived in the present study. However, they only found associations between this dimension and task-evoked brain activation, with no links to brain structural features. A plausible explanation is that their study used conventional brain structural features, which may be less sensitive than deviation scores for capturing structural correlates of psychopathology. Still, other factors, such as differing age ranges and multimodal inputs that may have dominated the sCCA solution, could also account for this inconsistency. Future studies could combine multimodal neuroimaging with brain deviation scores to improve effect sizes and generalizability of psychopathological dimensions.

Emotion dysregulation is a transdiagnostic clinical phenotype. Clinically, it is characterized by maladaptive emotional responses including high arousal, emotional lability, irritability, aggression, and temper outbursts, which resemble the symptom items identified in the present study ^40^. Prior work has linked emotion dysregulation in youth to reduced gray matter volume in the dorsolateral prefrontal and orbitofrontal cortices ^41,42^. However, gray matter volume reflects the combined contribution of surface area and thickness, which have distinct genetic bases and developmental trajectories ^43–45^. In this study, lower surface area of the middle frontal gyrus and orbitofrontal cortex was associated with more severe emotion dysregulation symptoms, whereas no significant correlations were detected for cortical thickness in these regions. Furthermore, reduced surface area in the superior temporal gyrus, insula, anterior cingulate gyrus, and posterior cingulate gyrus correlated with greater emotion dysregulation symptom severity. These brain regions are implicated in the recognition of vocal emotional cues ^46^, integration of diverse sensory inputs ^47,48^, conflict monitoring ^49^, rumination, and excessive self-referential processing ^50^. Collectively, our findings further suggest that surface area may represent a potential neurostructural substrate for emotional dysregulation in children and adolescents, which awaits further validation in larger independent cohorts.

In addition, we observed that brain regions showing abnormal cortical thickness were distinct from those with surface area, mainly including the inferior parietal lobule, precuneus, banks of the superior temporal sulcus, and postcentral gyrus. These regions have all been previously linked to aberrant emotional processing ^51–54^. Nevertheless, both the contribution magnitude and the number of brain regions involved were smaller for cortical thickness relative to surface area. This finding aligns with Blok et al. who showed that brain deviation scores of surface area exhibited more widespread and stronger associations with emotion dysregulation than cortical thickness ^55^. Furthermore, large-scale ENIGMA studies in children and adolescents have demonstrated that surface area yields stronger effect sizes than cortical thickness when associated with diverse internalizing and externalizing problems.

No significant associations between subcortical volumes and emotion dysregulation were observed in the present study. Findings regarding associations between emotion dysregulation and subcortical volumes remain inconsistent across child-adolescent samples. One previous study reported that reduced pallidus volume was linked to emotion dysregulation ^58^, whereas another investigation detected no significant subcortical correlates ^41^. These discrepancies may be attributable to limited sample size, differences in age distribution, and different measurement approaches for emotion dysregulation.

The present study identified four transdiagnostic psychopathological dimensions in the discovery dataset, including emotion dysregulation, externalizing behavior, eating pathology, and substance use and psychosis. The first three dimensions have been consistently documented in prior neuroimaging investigations ^9,10,12,59,60^, while the substance use and psychosis dimension represents a novel finding in the current study, consistent with existing evidence that substance use is a prominent risk factor for psychotic symptoms ^61,62^.

However, generalizability analyses revealed weak generalizability across the four aforementioned dimensions, with only emotion dysregulation showing a nominally significant correlation in the independent dataset. Emotion dysregulation is a broad transdiagnostic construct and a core component of the general psychopathology *p*-factor within the HiTOP framework ^63–65^. Its high explanatory variance allows sCCA to stably capture its latent patterns. By contrast, the remaining dimensions represent more specific phenotypes with limited individual variation, yielding less stable model weight estimation. Thus, larger samples are required to derive stable dimensional structures, as studies have shown more reliable weight estimates emerge across tens of thousands of participants ^66^. In addition, the independent sample in the present study consisted predominantly of typically-developing individuals, which may limit interindividual variability in psychiatric symptom severity.

In the present study, latent brain scores for emotion dysregulation showed significant associations with out-of-model emotion-dysregulation-related phenotypes, while no significant correlations were observed for non-emotion-dysregulation phenotypes. These findings suggest that latent brain scores, serving as a parsimonious composite index of brain deviation scores, capture the magnitude of emotion dysregulation specific brain abnormalities. Future work may adopt more sophisticated multivariate algorithms to capture nonlinear brain-behavior relationships and improve model interpretability ^67^.

The latent behavioral scores of emotion dysregulation increased with age in females but decreased with age in males, consistent with a recent longitudinal study ^68^. More importantly, females in the present study exhibited significantly higher latent brain scores relative to males. Prior work has identified female-specific brain functional signatures that predicts mood dysregulation symptoms at a two-year follow-up ^69^. Extending these observations, our study further demonstrates sex-divergent brain-deviation patterns for emotion dysregulation. Nevertheless, latent brain scores decreased with age across both sexes, suggesting that structural brain abnormality patterns attenuate over development.

The limitations of the present study are as follows. First, the psychopathological dimensions identified in this study showed weak brain-behavior correlations and limited generalizability, which collectively constrained their clinical and translational utility. Second, the brain-deviation patterns were derived from cross-sectional data and could not validate age-related effects. Future studies using large-sample, multimodal longitudinal cohorts are needed to identify robust psychopathological dimensions with larger effect sizes and examine their age-dependent developmental changes.

Neuroimaging-informed dimensional partitioning of psychopathology enables the identification of biologically meaningful transdiagnostic dimensions. Extending prior work, we show that brain deviation scores derived from normative modeling outperform conventional structural metrics in detecting transdiagnostic psychopathological dimensions in youth, offering new insights for constructing biologically informed dimensional frameworks in psychopathology research. We further found that cortical thickness and surface area brain deviation scores exhibited distinct regional distributions and weighting patterns in their contribution to emotion dysregulation. Finally, latent scores can act as a concise indicator of brain deviation scores by capturing dimension-specific brain abnormalities and improving the interpretability of brain deviation metrics.

## AUTHORSHIP CONTRIBUTIONS

Zheng Li: Methodology, Investigation, Data curation, Conceptualization, Writing – review & editing.

Bingqian Ren: Writing – review & editing.

Hao Wu: Writing – review & editing.

Zhidong Wang: Writing – review & editing.

Libin Zhang: Writing – review & editing.

## ACKNOWLEDGEMENTS

The authors are grateful to all participants for their contribution to this research.

## DATA AVAILABILITY

sMRI data from HBN and NKI-RS used in this study can be downloaded at. Access to behavioral data from HBN and NKI-RS requires a signed data-usage agreement (HBN: https://fcon_1000.projects.nitrc.org/indi/cmi_healthy_brain_network/Phenotypic.html; NKI-RS: https://rocklandsample.org/for-researchers).

Behavioral and brain weight coefficients for the emotion-dysregulation dimension identified via sCCA in the present study will be made publicly available upon publication.

## CODE AVAILABILITY

Our sCCA code was adapted from the open-source repository ^10^

(https://github.com/cedricx/sCCA/tree/master/sCCA/code/final). Custom scripts for the sCCA model and generalizability validation will be made publicly available upon publication.

