## Supplementary Information for "Brain deviation scores derived from normative modelling identify dimensions of psychopathology in children and adolescents"

1. **Supplementary Method;**
2. **Supplementary Figures 1-4;**
3. **Supplementary Tables 1-6;**

**Supplementary Method**

**Clinical diagnoses**

HBN adopted the computer‑assisted Schedule for Affective Disorders and Schizophrenia‑Children’s Version ( KSADS; Kaufman et al., 1997) for clinical psychiatric diagnoses. As a semi‑structured interview conducted by trained clinicians, the KSADS generates diagnostic information according to DSM‑5 criteria. To arrive at final diagnostic assignments, a panel of licensed clinicians synthesized interview results together with supplementary study‑derived records to reach group consensus.

Psychiatric assessments in NKI‑RS were conducted using the KSADS; participants turning 18 years of age during follow‑up were assessed with the Structured Clinical Interview for DSM‑Disorders‑NP ( SCID‑I/NP; First et al., 2002). Final consensus research diagnoses were derived by a panel of licensed clinicians following the best‑estimate method, synthesizing KSADS‑PL/SCID‑I/NP outputs together with all additional study‑collected information such as medical history and behavioral questionnaire data.

**Quality control and processing of sMRI data**

Inconsistent quality control (QC) criteria and data processing pipelines across independent cohorts represent a major confounding factor that impairs the reliability and cross-cohort comparability of neuroimaging studies. To address this methodological limitation, the present study adopted harmonized sMRI data from both discovery and independent datasets, which were uniformly quality-controlled and preprocessed by the Reproducible Brain Charts (RBC) consortium ^3^.

All sMRI scans underwent unified QC procedures, including visual inspection conducted by two to five trained raters who independently classified each scan as either "Pass" or "Fail". Specifically, scans receiving unanimous "Pass" ratings from all reviewers were assigned a final "Pass" label; scans with discrepant ratings (both "Pass" and "Fail") across raters were marked as artifact; and scans with unanimous "Fail" ratings were excluded from subsequent analyses. Only sMRI scans with a final "Pass" quality grade were included in the current study.

Following rigorous RBC-standardized quality assurance, uniform structural preprocessing was performed. Intensity inhomogeneity correction and skull stripping were implemented via the ANTs brain extraction pipeline embedded in sMRIPrep v0.7.1 ^4^. Whole-brain surface reconstruction was subsequently conducted using FreeSurfer v6.0.1 ^5^ to generate quantitative brain morphological features. Region-specific morphological metrics were derived based on standardized atlas-based segmentation. As shown in Figure 2A, regional cortical thickness and surface area were estimated using the Desikan–Killiany (DK; Desikan et al., 2006) atlas, while subcortical volumetric indices were segmented and quantified using the automatic segmentation (ASEG; Fischl et al., 2002) atlas. In total, 150 brain structural features were extracted: 136 regional cortical measures (68 regions × 2 metrics: cortical thickness and surface area) based on the Desikan–Killiany atlas, and 14 subcortical volume measures derived from the ASEG atlas, totaling 150 features.

**The sCCA model construction in the discovery cohort**

we implemented sCCA to identify brain patterns associated with CBCL items using the R package PMA (v. 1.2‑4). As a multivariate dimension‑reduction technique, sCCA maximizes correlation between linear combinations of two distinct variable sets. Sparsity at the feature level is enforced via $L_{1}$‑norm penalties imposed on loading vectors.

Let $\boldsymbol{X}_{n\times p}$ denote the matrix of brain deviation features and $\boldsymbol{Y}_{n\times q}$ denote the matrix of clinical‑symptom features, where n corresponds to the total number of participants, p is the count of brain features, and q denotes the number of clinical symptom indicators. All brain deviation scores and CBCL item scores were z-score standardized across participants to ensure equal variance across variables. As formulated in Equation (1), sCCA estimates loading vectors $\mathbf{u}$and $\mathbf{v}$ to maximize the canonical correlation between $\boldsymbol{X}\mathbf{u}$ and $\boldsymbol{Y}\mathbf{v}$.

Both loading vectors are constrained to have a unit$L_{2}$ norm for scale identifiability. Two separate $L_{1}$sparsity parameters $c_{1}$and $c_{2}$ are applied to the brain‑feature loading $\mathbf{u}$ and symptom‑feature loading $\mathbf{v}$, respectively. Critically, this unit‑$L_{2}$constraint serves purely to fix vector scale and does not introduce ridge‑type $L_{2}$regularization; sparsity arises exclusively from the $L_{1}$‑norm constraints.

$\begin{aligned} {argmax}_{u,v}\mathrm{Corr}\left( \boldsymbol{X}u,\boldsymbol{Y}v \right)\quad\text{s.t.}\left\{ \begin{matrix} \left\| u \right\|_{2}^{2}=1,\quad\left\| u \right\|_{1}\leq c_{1} \\ \left\| v \right\|_{2}^{2}=1,\quad\left\| v \right\|_{1}\leq c_{2} \end{matrix} \right. \end{aligned}$ (1)

Within the PMA package, sparsity parameters take values between 0 and 1: a value of 0 imposes maximal sparsity to select a minimal set of features, whereas a value of 1 produces fully dense loadings with all input features preserved. To select optimal $c_{1}$and $c_{2}$, we implemented a grid search with a step size of 0.1. Model parameters were optimized based on the averaged canonical correlation of the first canonical component across 10 independent resampling runs. For each iteration, two‑thirds of participants from the discovery dataset were randomly sampled.

Permutation testing was implemented to quantify the statistical significance for each canonical component. We kept the brain‑deviation matrix unchanged, while permuting row indices of the clinical‑symptom matrix to disrupt subject‑specific pairing between brain morphological deviation patterns and psychiatric symptom profiles. Holding regularization parameters fixed, we reran sCCA across 1000 permuted datasets to construct the null distribution of canonical correlation coefficients.

Permutation may induce arbitrary axis rotation or reflection, which can re‑order canonical components and invert the sign of loading weights. To resolve this misalignment, we matched canonical components obtained from each permuted dataset against those derived from the original data based on clinical‑symptom loading vectors. Empirical permutation p‑values were computed as the fraction of null‑distribution correlation coefficients exceeding the true canonical correlation observed in the non‑permuted dataset. This permutation strategy is well‑established in prior literature ^9,10^. Permuting participant labels importantly preserves the within‑dataset covariance structures of both clinical and neuroimaging measures. Finally, false‑discovery‑rate (FDR) correction was applied to empirical p‑values of the top four canonical components (based on the scree plot of covariance explained). The significance threshold was set to $p_{\text{FDR}}<0.05$.

We adopted a resampling framework with 1000 iterations to quantify the stability of features within each canonical component. At every iteration, two‑thirds of participants in the discovery cohort were randomly subsampled; the remaining one‑third of observations were then generated via resampling with replacement from this selected subset. The distribution of weights for each feature derived from repeated resampling was used to construct its corresponding confidence interval. Following the matching strategy used for permutation analyses, canonical components estimated on each resampled dataset were aligned against components from the original data, permitting valid cross‑decomposition comparisons. A feature was defined as robust and stable when its resampling‑based confidence interval excluded zero. We applied a 95 % confidence interval (CI) threshold for clinical symptom variables. By contrast, we adopted a stricter 99 % CI threshold for brain‑deviation features owing to the higher dimensionality of brain measures.

**Measurement of phenotypes used for specificity validation**

To verify the dimensional specificity of the emotion dysregulation dimension, partial correlations were computed between latent scores (brain and behavior separately) of each psychopathological dimension and out‑of‑model phenotypes, with the latent scores of the remaining three dimensions included as covariates.

Phenotype selection was guided by the psychopathological dimensions identified in the present study. Specifically, emotion dysregulation is linked to irritability ^11^. Callous‑unemotional traits are associated with severe and chronic externalizing behaviors ^12^. Eating‑related pathology specifically correlates with BMI ^13^. Cocaine use represents a prevalent form of substance use among adolescents ^14^.

We hypothesized that if the generalizable emotion dysregulation dimension identified via sCCA exhibits true specificity, its corresponding latent brain scores would show robust associations with irritability, anxiety, and depression but weaker correlations with the remaining behavioral phenotypes.

Irritability was assessed with the Affective Reactivity Index (ARI)^15^. The ARI consists of six items rated on a 3‑point scale. Summing these six items produces a total score, with higher values indicating greater severity of persistent irritability. The ARI exhibits robust psychometric properties, including excellent internal consistency (Cronbach’s α = 0.89–0.92). In the discovery dataset, 1,075 participants completed the parent‑report ARI. Partial correlation between the latent scores of emotional‑dysregulation and the parent‑report ARI total score was computed within this sample.

Callous‑unemotional (CU) traits were measured with the Inventory of Callous‑Unemotional Traits (ICU) ^16^. The parent‑report version was used in the current study. The ICU demonstrates favorable psychometric properties and good internal consistency (Cronbach’s α ≈ 0.80). In the discovery dataset, 998 participants completed the parent‑report ICU. Partial correlation was calculated between latent scores of the emotion dysregulation dimension and parent‑report ICU total scores within this subsample.

Body mass index (BMI, kg/m²) was calculated from the bioelectric impedance analysis. In the discovery dataset, 830 participants completed the assessment. Partial correlation was calculated between latent scores of the emotion dysregulation dimension and BMI within this subsample.

Cocaine‑related substance involvement was assessed using the cocaine involvement score derived from the NIDA‑Modified ASSIST. This screening instrument queries lifetime substance exposure, as well as past‑3‑month frequency of substance use, substance‑use craving, functional impairment attributable to substance use, and interpersonal concerns regarding participants’ substance‑using behaviors. In the discovery dataset, 32 participants had valid data for the NIDA‑Modified ASSIST. Partial correlations were calculated between latent scores of the emotion dysregulation dimension and cocaine involvement score within this subsample.

**Supplementary Figures**

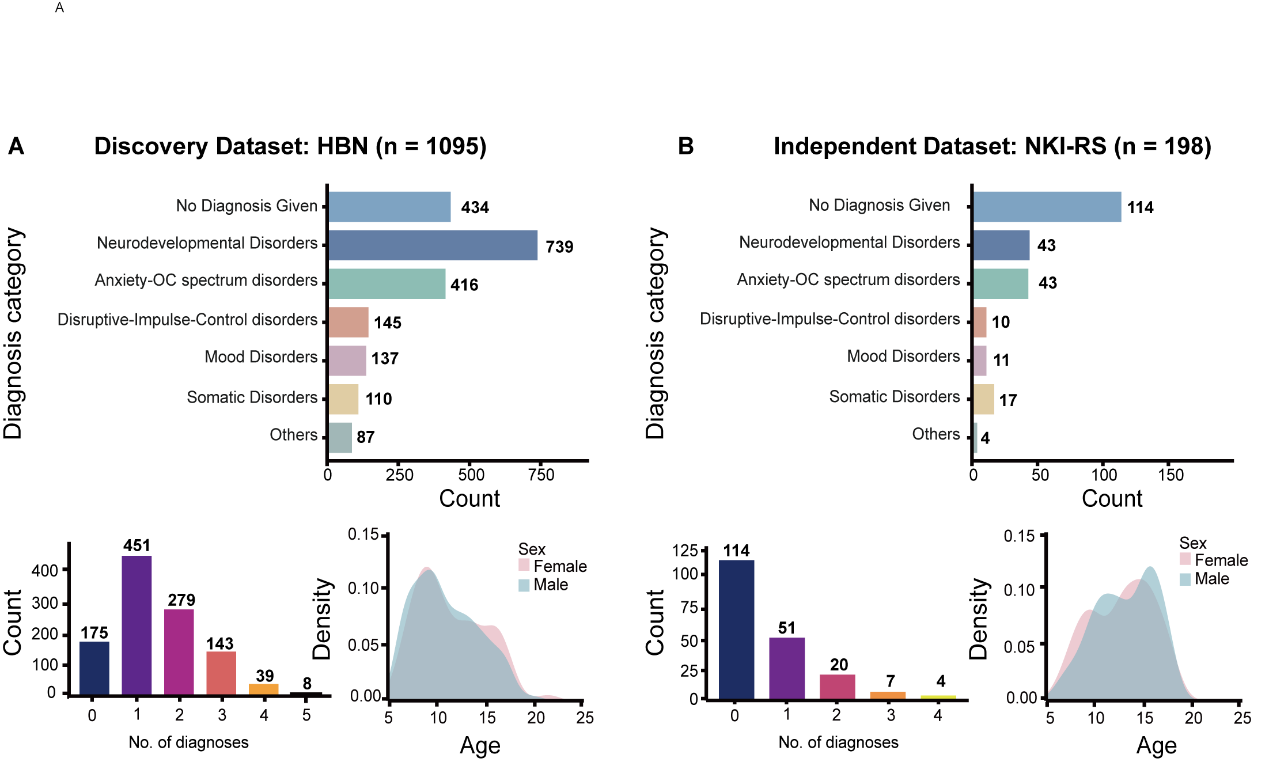

**Supplementary Figure 1**. Sample characteristics of the discovery and independent datasets. (A) HBN discovery dataset. (B) NKI‑RS independent dataset. For both datasets, the upper bar plot shows the distribution of psychiatric diagnosis count; the lower‑left plot shows comorbidity count; the lower‑right plot shows age and sex distribution. HBN: Healthy Brain Network; NKI‑RS: NKI‑Rockland Sample. Notes: two NKI‑RS participants lacked categorical diagnostic information, leaving 196 with available diagnostic and comorbidity data.

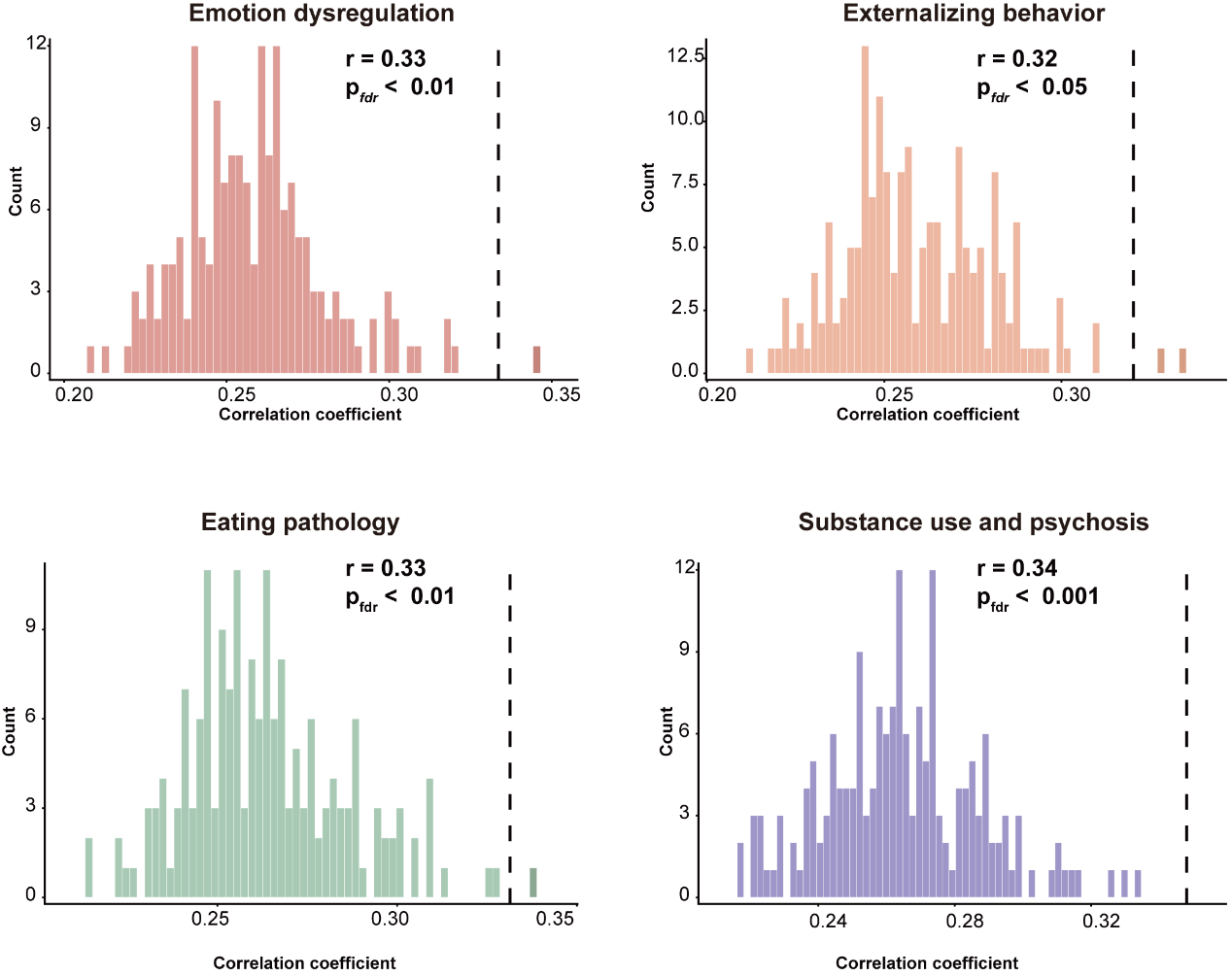
 **Supplementary Figure 2**. Permutation‑test results for canonical variables of brain deviation-based sCCA model.

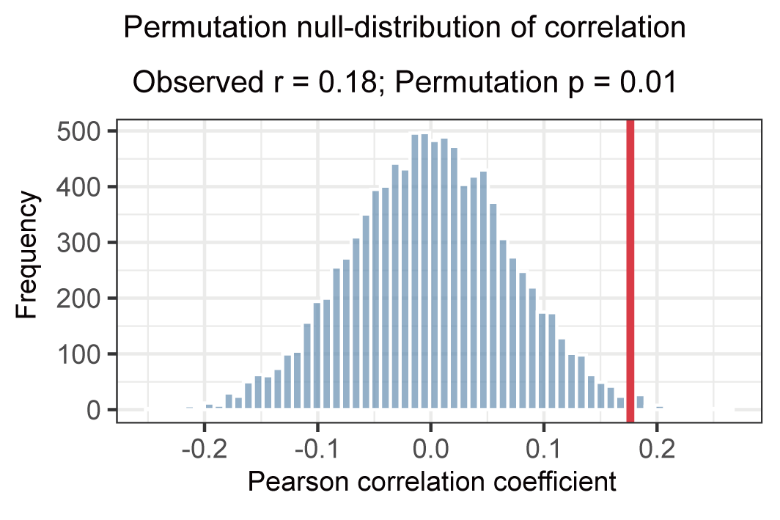

**Supplementary Figure 3.** Permutation null‑distribution for the Pearson correlation between latent brain and behavior of emotion dysregulation in the independent dataset. The histogram shows the distribution of correlation coefficients generated under the null hypothesis by 10 000 permutations. The solid red vertical line denotes the observed correlation coefficient ($r=0.169$). The permutation‑derived p‑value was 0.016.

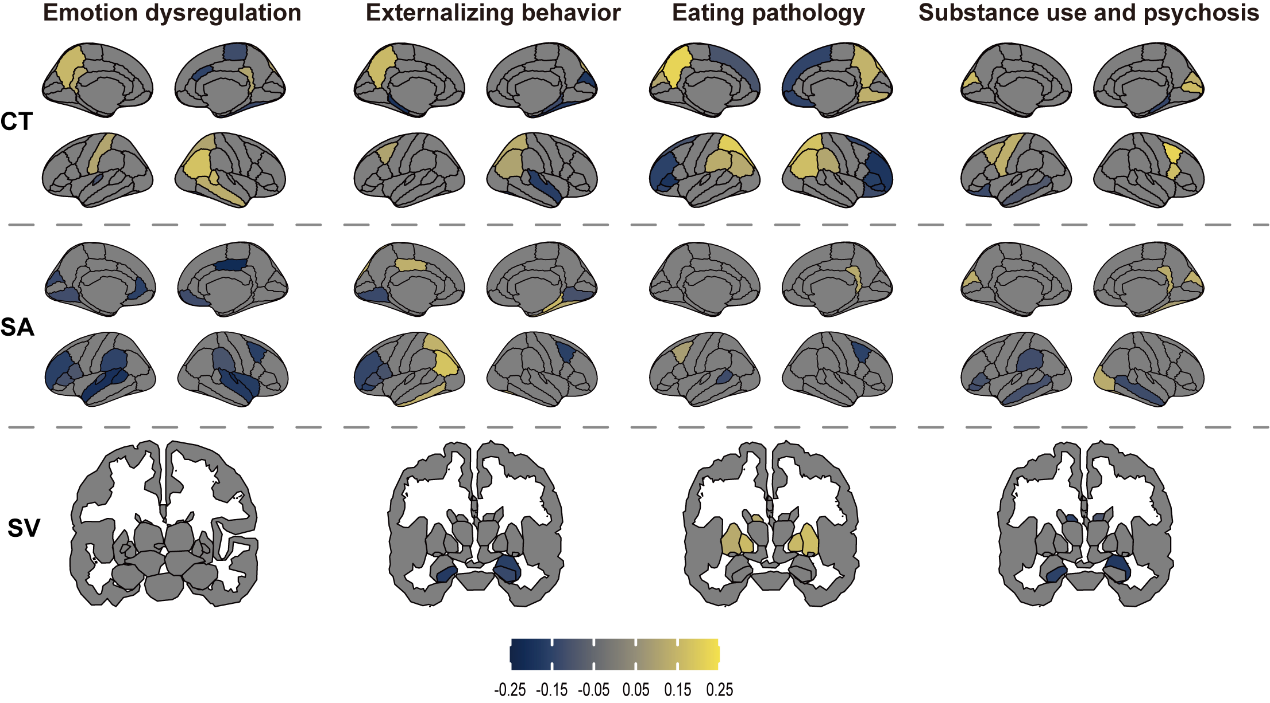

**Supplementary Figure 4**. Distinct brain deviation patterns across canonical variables

**Supplementary Tables**

**Supplementary Table 1.** Count of Participants by Categorical Diagnostic Status in HBN

| **category** | **DSM diagnosis** | **count** | **sum of category** |
| --- | --- | --- | --- |
| Neurodevelopmental Disorders | Neurodevelopmental Disorders | 739 | 739 |
| Mood Disorders | Depressive Disorders | 132 | 137 |
|  | Bipolar and Related Disorders | 5 |  |
| Anxiety and OC Spectrum Disorders | Anxiety Disorders | 364 | 416 |
|  | Obsessive Compulsive and Related Disorders | 52 |  |
| Disruptive, Impulse-Control and Conduct Disorders | Disruptive, Impulse Control and Conduct Disorders | 145 | 145 |
| Somatic Disorders | Elimination Disorders | 96 | 110 |
|  | Feeding and Eating Disorders | 12 |  |
|  | Sleep-Wake Disorders | 1 |  |
|  | Somatic Symptom and Related Disorders | 1 |  |
| Others | Trauma and Stressor Related Disorders | 55 | 1 |
|  | Substance Related and Addictive Disorders | 13 | 87 |
|  | Schizophrenia Spectrum and other Psychotic Disorders | 5 |  |
|  | Neurocognitive Disorders | 1 |  |
|  | Other Conditions That May Be a Focus of Clinical Attention | 13 |  |
| No Diagnosis Given | No Diagnosis Given | 434 | 434 |

**Supplementary Table 2.** Count of Participants by Categorical Diagnostic Status in NKI‑RS

| **category** | **DSM diagnosis** | **count** | **sum of category** |
| --- | --- | --- | --- |
| Neurodevelopmental Disorders | Attention-Deficit/Hyperactivity Disorder, Predominantly Inattentive Type | 13 | 43 |
|  | Attention-Deficit/Hyperactivity Disorder NOS | 11 |  |
|  | Attention-Deficit/Hyperactivity Disorder, Combined Type | 8 |  |
|  | Attention-Deficit/Hyperactivity Disorder, Predominantly Hyperactive-Impulsive Type | 1 |  |
|  | Chronic Motor or Vocal Tic Disorder | 5 |  |
|  | Transient Tic Disorder | 3 |  |
|  | Tic Disorder NOS | 1 |  |
|  | Tourette's Disorder | 1 |  |
| Mood Disorders | Major Depressive Disorder, Single Episode, In Full Remission | 4 | 11 |
|  | Major Depressive Disorder, Recurrent, In Full Remission | 2 |  |
|  | Major Depressive Disorder, Single Episode, Unspecified | 2 |  |
|  | Depressive Disorder NOS | 1 |  |
|  | Dysthymic Disorder | 1 |  |
|  | Major Depressive Disorder, Single Episode, Severe Without Psychotic Features | 1 |  |
| Anxiety and OC Spectrum Disorders | Generalized Anxiety Disorder | 17 | 43 |
|  | Specific Phobia | 9 |  |
|  | Separation Anxiety Disorder | 5 |  |
|  | Obsessive-Compulsive Disorder | 4 |  |
|  | Social Phobia | 3 |  |
|  | Anxiety Disorder NOS | 2 |  |
|  | Panic Disorder Without Agoraphobia | 2 |  |
|  | Panic Disorder With Agoraphobia | 1 |  |
| Disruptive, Impulse-Control and Conduct Disorders | Oppositional Defiant Disorder | 9 | 10 |
|  | Disruptive Behavior Disorder NOS | 1 |  |
| Somatic Disorders | Enuresis (Not Due to a General Medical Condition) | 15 | 17 |
|  | Encopresis, Without Constipation and Overflow Incontinence | 2 |  |
| Others | Cannabis Abuse | 3 | 4 |
|  | Posttraumatic Stress Disorder | 1 |  |
| No Diagnosis Given | No Diagnosis Given | 114 | 114 |

**Supplementary Table 3**. Order of CBCL items used for sCCA input. The first column indicates the order of CBCL items within the behavioral matrix used as input for sCCA. The second column corresponds to the original item number of the CBCL scale. The third column lists the verbatim CBCL item descriptions.

| **behavioral matrix order** | **CBCL item order** | **question** |
| --- | --- | --- |
| 1 | 1 | Acts too young for his/her age |
| 2 | 2 | Drinks alcohol without parents' approval (please describe) |
| 3 | 3 | Argues a lot |
| 4 | 4 | Fails to finish things he/she starts |
| 5 | 5 | There is very little he/she enjoys |
| 6 | 6 | Bowel movements outside toilet |
| 7 | 7 | Bragging, boasting |
| 8 | 8 | Can't concentrate, can't pay attention for long |
| 9 | 9 | Can't get his/her mind off certain thoughts; obsessions (please describe) |
| 10 | 10 | Can't sit still, restless or hyperactive |
| 11 | 100 | Trouble sleeping (please describe) |
| 12 | 101 | Truancy, skips school |
| 13 | 102 | Underactive, slow moving, or lacks energy |
| 14 | 103 | Unhappy, sad, or depressed |
| 15 | 104 | Unusually loud |
| 16 | 105 | Uses drugs for nonmedical purposes (don't include alcohol or tobacco) (please describe) |
| 17 | 106 | Vandalism |
| 18 | 107 | Wets self during the day |
| 19 | 108 | Wets the bed |
| 20 | 109 | Whining |
| 21 | 11 | Clings to adults or too dependent |
| 22 | 110 | Wishes to be of opposite sex |
| 23 | 111 | Withdrawn, doesn't get inolved with others |
| 24 | 112 | Worries |
| 25 | 12 | Complains of loneliness |
| 26 | 13 | Confused or seems to be in a fog |
| 27 | 14 | Cries a lot |
| 28 | 15 | Cruel to animals |
| 29 | 16 | Cruelty, bullying, or meanness to others |
| 30 | 17 | Daydreams or gets lost in his/her thoughts |
| 31 | 18 | Deliberately harms self or attempts suicide |
| 32 | 19 | Demands a lot of attention |
| 33 | 20 | Destroys his/her own things |
| 34 | 21 | Destroys things belonging to his/her family or others |
| 35 | 22 | Disobedient at home |
| 36 | 23 | Disobedient at school |
| 37 | 24 | Doesn't eat well |
| 38 | 25 | Doesn't get along well with other kids |
| 39 | 26 | Doesn't seem to feel guilty after misbehaving |
| 40 | 27 | Easily jealous |
| 41 | 28 | Breaks rules at home, school, or elsewhere |
| 42 | 29 | Fears certain animals, situations, or places, other than school (please describe) |
| 43 | 30 | Fears going to school |
| 44 | 31 | Fears he/she might think or do something bad |
| 45 | 32 | Feels he/she has to be perfect |
| 46 | 33 | Feels or complains that no one loves him/her |
| 47 | 34 | Feels others are out to get him/her |
| 48 | 35 | Feels worthless or inferior |
| 49 | 36 | Gets hurt a lot, accident-prone |
| 50 | 37 | Gets in many fights |
| 51 | 38 | Gets teased a lot |
| 52 | 39 | Hangs around with others who get in trouble |
| 53 | 40 | Hears sounds or voices that aren't there (please describe) |
| 54 | 42 | Would rather be alone than with others |
| 55 | 43 | Lying or cheating |
| 56 | 44 | Bites fingernails |
| 57 | 45 | Nervous, highstrung, or tense |
| 58 | 46 | Nervous movements or twitching (please describe) |
| 59 | 47 | Nightmares |
| 60 | 48 | Not liked by other kids |
| 61 | 49 | Constipated, doesn't move bowels |
| 62 | 50 | Too fearful or anxious |
| 63 | 51 | Feels dizzy or lightheaded |
| 64 | 52 | Feels too guilty |
| 65 | 53 | Overeating |
| 66 | 54 | Overtired without good reason |
| 67 | 55 | Overweight |
| 68 | 56A | Aches or pains (not stomach or headaches) |
| 69 | 56B | Headaches |
| 70 | 56C | Nausea, feels sick |
| 71 | 56D | Problems with eyes (not if corrected by glasses (please describe) |
| 72 | 56E | Rashes or other skin problems |
| 73 | 56F | Stomachaches |
| 74 | 56G | Vomiting, throwing up |
| 75 | 57 | Physically attacks people |
| 76 | 58 | Picks nose, skin, or other parts of body (please describe) |
| 77 | 59 | Plays with own sex parts in public |
| 78 | 60 | Plays with own sex parts too much |
| 79 | 61 | Poor school work |
| 80 | 62 | Poorly coordinated or clumsy |
| 81 | 63 | Prefers being with older kids |
| 82 | 64 | Prefers being with younger kids |
| 83 | 65 | Refuses to talk |
| 84 | 66 | Repeats certain acts over and over; compulsions (please describe) |
| 85 | 67 | Runs away from home |
| 86 | 68 | Screams a lot |
| 87 | 69 | Secretive, keeps things to self |
| 88 | 70 | Sees things that aren't there (please describe) |
| 89 | 71 | Self-conscious or easily embarrassed |
| 90 | 72 | Sets fires |
| 91 | 73 | Sexual problems (please describe) |
| 92 | 74 | Showing off or clowning |
| 93 | 75 | Too shy or timid |
| 94 | 76 | Sleeps less than most kids |
| 95 | 77 | Sleeps more than most kids during day and/or night (please describe) |
| 96 | 78 | Inattentive or easily distracted |
| 97 | 79 | Speech problem (please describe) |
| 98 | 80 | Stares blankly |
| 99 | 81 | Steals at home |
| 100 | 82 | Steals outside the home |
| 101 | 83 | Stores up too many things he/she doesn't need (please describe) |
| 102 | 84 | Strange behavior (please describe) |
| 103 | 85 | Strange ideas (please describe) |
| 104 | 86 | Stubborn, sullen, or irritable |
| 105 | 87 | Sudden changes in mood or feelings |
| 106 | 88 | Sulks a lot |
| 107 | 89 | Suspicious |
| 108 | 90 | Swearing or obscene language |
| 109 | 91 | Talks about killing self |
| 110 | 92 | Talks or walks in sleep (please describe) |
| 111 | 93 | Talks too much |
| 112 | 94 | Teases a lot |
| 113 | 95 | Temper tantrums or hot temper |
| 114 | 96 | Thinks about sex too much |
| 115 | 97 | Threatens people |
| 116 | 98 | Thumb-sucking |
| 117 | 99 | Smokes, chews, or sniffs tobacco |

**Supplementary Table 4**. Variable order within the brain matrix for sCCA input. Column 1 shows the variable order in the brain matrix fed into sCCA. Column 2 indicates hemisphere information (left/right). Column 3 denotes the type of brain morphological metric. Column 4 lists the brain region name. The matrices for brain deviation scores and brain structural features both followed this ordering.

| **brain matrix order** | **hemi** | **morphological metrics** | **region** |
| --- | --- | --- | --- |
| 1 | left | thickness | banks of superior temporal sulcus |
| 2 | left | thickness | caudal anterior cingulate |
| 3 | left | thickness | caudal middle frontal |
| 4 | left | thickness | cuneus |
| 5 | left | thickness | entorhinal |
| 6 | left | thickness | fusiform |
| 7 | left | thickness | inferior parietal |
| 8 | left | thickness | inferior temporal |
| 9 | left | thickness | isthmus cingulate |
| 10 | left | thickness | lateral occipital |
| 11 | left | thickness | lateral orbitofrontal |
| 12 | left | thickness | lingual |
| 13 | left | thickness | medial orbitofrontal |
| 14 | left | thickness | middle temporal |
| 15 | left | thickness | parahippocampal |
| 16 | left | thickness | paracentral |
| 17 | left | thickness | pars opercularis |
| 18 | left | thickness | pars orbitalis |
| 19 | left | thickness | pars triangularis |
| 20 | left | thickness | pericalcarine |
| 21 | left | thickness | postcentral |
| 22 | left | thickness | posterior cingulate |
| 23 | left | thickness | precentral |
| 24 | left | thickness | precuneus |
| 25 | left | thickness | rostral anterior cingulate |
| 26 | left | thickness | rostral middle frontal |
| 27 | left | thickness | superior frontal |
| 28 | left | thickness | superior parietal |
| 29 | left | thickness | superior temporal |
| 30 | left | thickness | supramarginal |
| 31 | left | thickness | frontal pole |
| 32 | left | thickness | temporal pole |
| 33 | left | thickness | transverse temporal |
| 34 | left | thickness | insula |
| 35 | right | thickness | banks of superior temporal sulcus |
| 36 | right | thickness | caudal anterior cingulate |
| 37 | right | thickness | caudal middle frontal |
| 38 | right | thickness | cuneus |
| 39 | right | thickness | entorhinal |
| 40 | right | thickness | fusiform |
| 41 | right | thickness | inferior parietal |
| 42 | right | thickness | inferior temporal |
| 43 | right | thickness | isthmus cingulate |
| 44 | right | thickness | lateral occipital |
| 45 | right | thickness | lateral orbitofrontal |
| 46 | right | thickness | lingual |
| 47 | right | thickness | medial orbitofrontal |
| 48 | right | thickness | middle temporal |
| 49 | right | thickness | parahippocampal |
| 50 | right | thickness | paracentral |
| 51 | right | thickness | pars opercularis |
| 52 | right | thickness | pars orbitalis |
| 53 | right | thickness | pars triangularis |
| 54 | right | thickness | pericalcarine |
| 55 | right | thickness | postcentral |
| 56 | right | thickness | posterior cingulate |
| 57 | right | thickness | precentral |
| 58 | right | thickness | precuneus |
| 59 | right | thickness | rostral anterior cingulate |
| 60 | right | thickness | rostral middle frontal |
| 61 | right | thickness | superior frontal |
| 62 | right | thickness | superior parietal |
| 63 | right | thickness | superior temporal |
| 64 | right | thickness | supramarginal |
| 65 | right | thickness | frontal pole |
| 66 | right | thickness | temporal pole |
| 67 | right | thickness | transverse temporal |
| 68 | right | thickness | insula |
| 69 | left | surface area | banks of superior temporal sulcus |
| 70 | left | surface area | caudal anterior cingulate |
| 71 | left | surface area | caudal middle frontal |
| 72 | left | surface area | cuneus |
| 73 | left | surface area | entorhinal |
| 74 | left | surface area | fusiform |
| 75 | left | surface area | inferior parietal |
| 76 | left | surface area | inferior temporal |
| 77 | left | surface area | isthmus cingulate |
| 78 | left | surface area | lateral occipital |
| 79 | left | surface area | lateral orbitofrontal |
| 80 | left | surface area | lingual |
| 81 | left | surface area | medial orbitofrontal |
| 82 | left | surface area | middle temporal |
| 83 | left | surface area | parahippocampal |
| 84 | left | surface area | paracentral |
| 85 | left | surface area | pars opercularis |
| 86 | left | surface area | pars orbitalis |
| 87 | left | surface area | pars triangularis |
| 88 | left | surface area | pericalcarine |
| 89 | left | surface area | postcentral |
| 90 | left | surface area | posterior cingulate |
| 91 | left | surface area | precentral |
| 92 | left | surface area | precuneus |
| 93 | left | surface area | rostral anterior cingulate |
| 94 | left | surface area | rostral middle frontal |
| 95 | left | surface area | superior frontal |
| 96 | left | surface area | superior parietal |
| 97 | left | surface area | superior temporal |
| 98 | left | surface area | supramarginal |
| 99 | left | surface area | frontal pole |
| 100 | left | surface area | temporal pole |
| 101 | left | surface area | transverse temporal |
| 102 | left | surface area | insula |
| 103 | right | surface area | banks of superior temporal sulcus |
| 104 | right | surface area | caudal anterior cingulate |
| 105 | right | surface area | caudal middle frontal |
| 106 | right | surface area | cuneus |
| 107 | right | surface area | entorhinal |
| 108 | right | surface area | fusiform |
| 109 | right | surface area | inferior parietal |
| 110 | right | surface area | inferior temporal |
| 111 | right | surface area | isthmus cingulate |
| 112 | right | surface area | lateral occipital |
| 113 | right | surface area | lateral orbitofrontal |
| 114 | right | surface area | lingual |
| 115 | right | surface area | medial orbitofrontal |
| 116 | right | surface area | middle temporal |
| 117 | right | surface area | parahippocampal |
| 118 | right | surface area | paracentral |
| 119 | right | surface area | pars opercularis |
| 120 | right | surface area | pars orbitalis |
| 121 | right | surface area | pars triangularis |
| 122 | right | surface area | pericalcarine |
| 123 | right | surface area | postcentral |
| 124 | right | surface area | posterior cingulate |
| 125 | right | surface area | precentral |
| 126 | right | surface area | precuneus |
| 127 | right | surface area | rostral anterior cingulate |
| 128 | right | surface area | rostral middle frontal |
| 129 | right | surface area | superior frontal |
| 130 | right | surface area | superior parietal |
| 131 | right | surface area | superior temporal |
| 132 | right | surface area | supramarginal |
| 133 | right | surface area | frontal pole |
| 134 | right | surface area | temporal pole |
| 135 | right | surface area | transverse temporal |
| 136 | right | surface area | insula |
| 137 | left | subcortical volume | Thalamus |
| 138 | right | subcortical volume | Thalamus |
| 139 | left | subcortical volume | Caudate |
| 140 | right | subcortical volume | Caudate |
| 141 | left | subcortical volume | Putamen |
| 142 | right | subcortical volume | Putamen |
| 143 | left | subcortical volume | Pallidum |
| 144 | right | subcortical volume | Pallidum |
| 145 | left | subcortical volume | Hippocampus |
| 146 | right | subcortical volume | Hippocampus |
| 147 | left | subcortical volume | Amygdala |
| 148 | right | subcortical volume | Amygdala |
| 149 | left | subcortical volume | accumbens area |
| 150 | right | subcortical volume | accumbens area |

**Supplementary Table 5**. Variance explained by canonical variables derived from sCCA using brain deviation scores

| **canonical variable** | **variance explained** | **Cumulative explained variance** |
| --- | --- | --- |
| 1 | 0.12 | 12% |
| 2 | 0.07 | 19% |
| 3 | 0.06 | 25% |
| 4 | 0.04 | 29% |
| 5 | 0.04 | 33% |
| 6 | 0.03 | 36% |
| 7 | 0.03 | 39% |
| 8 | 0.03 | 42% |
| 9 | 0.02 | 44% |
| 10 | 0.03 | 47% |
| 11 | 0.02 | 49% |
| 12 | 0.02 | 51% |
| 13 | 0.02 | 53% |
| 14 | 0.02 | 55% |
| 15 | 0.02 | 57% |
| 16 | 0.02 | 59% |
| 17 | 0.02 | 61% |
| 18 | 0.02 | 63% |
| 19 | 0.02 | 65% |
| 20 | 0.01 | 66% |
| 21 | 0.01 | 67% |
| 22 | 0.01 | 68% |
| 23 | 0.01 | 69% |
| 24 | 0.01 | 70% |
| 25 | 0.01 | 71% |
| 26 | 0.01 | 72% |
| 27 | 0.01 | 73% |
| 28 | 0.01 | 74% |
| 29 | 0.01 | 75% |
| 30 | 0.01 | 76% |
| 31 | 0.01 | 77% |
| 32 | 0.01 | 78% |
| 33 | 0.01 | 79% |
| 34 | 0.01 | 80% |
| 35 | 0.01 | 81% |
| 36 | 0.01 | 82% |
| 37 | 0.01 | 83% |
| 38 | 0.01 | 84% |
| 39 | 0.01 | 85% |
| 40 | 0.01 | 86% |
| 41 | 0.01 | 87% |
| 42 | 0.01 | 88% |
| 43 | 0.01 | 89% |
| 44 | 0.01 | 90% |
| 45 | 0.01 | 91% |
| 46 | 0.00 | 91% |
| 47 | 0.00 | 91% |
| 48 | 0.00 | 91% |
| 49 | 0.00 | 91% |
| 50 | 0.01 | 92% |
| 51 | 0.00 | 92% |
| 52 | 0.00 | 92% |
| 53 | 0.00 | 92% |
| 54 | 0.00 | 92% |
| 55 | 0.00 | 92% |
| 56 | 0.00 | 92% |
| 57 | 0.00 | 92% |
| 58 | 0.00 | 92% |
| 59 | 0.00 | 92% |
| 60 | 0.00 | 92% |
| 61 | 0.00 | 92% |
| 62 | 0.00 | 92% |
| 63 | 0.00 | 92% |
| 64 | 0.00 | 92% |
| 65 | 0.00 | 92% |
| 66 | 0.00 | 92% |
| 67 | 0.00 | 92% |
| 68 | 0.00 | 92% |
| 69 | 0.00 | 92% |
| 70 | 0.00 | 92% |
| 71 | 0.00 | 92% |
| 72 | 0.00 | 92% |
| 73 | 0.00 | 92% |
| 74 | 0.00 | 92% |
| 75 | 0.00 | 92% |
| 76 | 0.00 | 92% |
| 77 | 0.00 | 92% |
| 78 | 0.00 | 92% |
| 79 | 0.00 | 92% |
| 80 | 0.00 | 92% |
| 81 | 0.00 | 92% |
| 82 | 0.00 | 92% |
| 83 | 0.00 | 92% |
| 84 | 0.00 | 92% |
| 85 | 0.00 | 92% |
| 86 | 0.00 | 92% |
| 87 | 0.00 | 92% |
| 88 | 0.00 | 92% |
| 89 | 0.00 | 92% |
| 90 | 0.00 | 92% |
| 91 | 0.00 | 92% |
| 92 | 0.00 | 92% |
| 93 | 0.00 | 92% |
| 94 | 0.00 | 92% |
| 95 | 0.00 | 92% |
| 96 | 0.00 | 92% |
| 97 | 0.00 | 92% |
| 98 | 0.00 | 92% |
| 99 | 0.00 | 92% |
| 100 | 0.00 | 92% |
| 101 | 0.00 | 92% |
| 102 | 0.00 | 92% |
| 103 | 0.00 | 92% |
| 104 | 0.00 | 92% |
| 105 | 0.00 | 92% |
| 106 | 0.00 | 92% |
| 107 | 0.00 | 92% |
| 108 | 0.00 | 92% |
| 109 | 0.00 | 92% |
| 110 | 0.00 | 92% |
| 111 | 0.00 | 92% |
| 112 | 0.00 | 92% |
| 113 | 0.00 | 92% |
| 114 | 0.00 | 92% |
| 115 | 0.00 | 92% |
| 116 | 0.00 | 92% |
| 117 | 0.00 | 92% |

**Supplementary Table 6**. Variance explained by canonical variables derived from sCCA using conventional brain structural features

| **canonical variable** | **variance explained** | **Cumulative explained variance** |
| --- | --- | --- |
| 1 | 0.03 | 3% |
| 2 | 0.02 | 5% |
| 3 | 0.03 | 8% |
| 4 | 0.02 | 10% |
| 5 | 0.02 | 12% |
| 6 | 0.02 | 14% |
| 7 | 0.02 | 16% |
| 8 | 0.02 | 18% |
| 9 | 0.02 | 19% |
| 10 | 0.02 | 21% |
| 11 | 0.02 | 23% |
| 12 | 0.02 | 24% |
| 13 | 0.02 | 26% |
| 14 | 0.02 | 28% |
| 15 | 0.02 | 29% |
| 16 | 0.01 | 31% |
| 17 | 0.01 | 32% |
| 18 | 0.01 | 34% |
| 19 | 0.01 | 35% |
| 20 | 0.00 | 35% |
| 21 | 0.01 | 36% |
| 22 | 0.01 | 37% |
| 23 | 0.01 | 38% |
| 24 | 0.01 | 40% |
| 25 | 0.01 | 41% |
| 26 | 0.01 | 42% |
| 27 | 0.01 | 43% |
| 28 | 0.01 | 44% |
| 29 | 0.02 | 46% |
| 30 | 0.01 | 47% |
| 31 | 0.01 | 49% |
| 32 | 0.01 | 50% |
| 33 | 0.00 | 50% |
| 34 | 0.01 | 51% |
| 35 | 0.01 | 52% |
| 36 | 0.01 | 53% |
| 37 | 0.01 | 55% |
| 38 | 0.00 | 55% |
| 39 | 0.01 | 56% |
| 40 | 0.01 | 57% |
| 41 | 0.01 | 58% |
| 42 | 0.01 | 59% |
| 43 | 0.00 | 59% |
| 44 | 0.01 | 60% |
| 45 | 0.01 | 61% |
| 46 | 0.01 | 61% |
| 47 | 0.01 | 62% |
| 48 | 0.01 | 63% |
| 49 | 0.01 | 64% |
| 50 | 0.01 | 65% |
| 51 | 0.01 | 66% |
| 52 | 0.01 | 67% |
| 53 | 0.01 | 68% |
| 54 | 0.00 | 68% |
| 55 | 0.00 | 68% |
| 56 | 0.01 | 69% |
| 57 | 0.01 | 69% |
| 58 | 0.01 | 70% |
| 59 | 0.00 | 70% |
| 60 | 0.01 | 71% |
| 61 | 0.01 | 71% |
| 62 | 0.00 | 72% |
| 63 | 0.01 | 72% |
| 64 | 0.01 | 73% |
| 65 | 0.01 | 74% |
| 66 | 0.01 | 75% |
| 67 | 0.01 | 75% |
| 68 | 0.01 | 76% |
| 69 | 0.01 | 77% |
| 70 | 0.01 | 77% |
| 71 | 0.01 | 78% |
| 72 | 0.01 | 79% |
| 73 | 0.01 | 80% |
| 74 | 0.01 | 81% |
| 75 | 0.01 | 82% |
| 76 | 0.00 | 82% |
| 77 | 0.00 | 82% |
| 78 | 0.01 | 83% |
| 79 | 0.00 | 83% |
| 80 | 0.01 | 84% |
| 81 | 0.01 | 85% |
| 82 | 0.00 | 85% |
| 83 | 0.01 | 86% |
| 84 | 0.00 | 86% |
| 85 | 0.01 | 87% |
| 86 | 0.00 | 87% |
| 87 | 0.00 | 87% |
| 88 | 0.01 | 88% |
| 89 | 0.01 | 89% |
| 90 | 0.01 | 89% |
| 91 | 0.00 | 89% |
| 92 | 0.00 | 89% |
| 93 | 0.00 | 89% |
| 94 | 0.00 | 90% |
| 95 | 0.00 | 90% |
| 96 | 0.01 | 91% |
| 97 | 0.01 | 91% |
| 98 | 0.00 | 92% |
| 99 | 0.01 | 92% |
| 100 | 0.00 | 93% |
| 101 | 0.00 | 93% |
| 102 | 0.01 | 94% |
| 103 | 0.00 | 94% |
| 104 | 0.01 | 94% |
| 105 | 0.01 | 95% |
| 106 | 0.01 | 96% |
| 107 | 0.01 | 96% |
| 108 | 0.01 | 97% |
| 109 | 0.00 | 97% |
| 110 | 0.01 | 98% |
| 111 | 0.00 | 98% |
| 112 | 0.00 | 98% |
| 113 | 0.01 | 99% |
| 114 | 0.01 | 99% |
| 115 | 0.00 | 99% |
| 116 | 0.00 | 100% |
| 117 | 0.00 | 100% |
